# The Step-Wise C-Truncation and Transport of ESyt3 to Lipid Droplets Reveals a Mother Primordial Cisterna

**DOI:** 10.1101/2020.07.17.209122

**Authors:** Vasiliki Lalioti, Galina V. Beznoussenko, Alexander A. Mironov, Ignacio V. Sandoval

## Abstract

Extended synaptotagmins (E-Syts) are endoplasmic reticulum (ER) proteins consisting of an SMP domain and multiple C2 domains that bind phospholipids and Ca^2+^. E-Syts create contact junctions between the ER and plasma membrane to facilitate lipid exchange. During adipocyte differentiation, the proteasome-based removal of the C2C domain results in targeting of E-Syt3 to the primordial cisterna, a previously undescribed giant annular organelle that mothers the LDs. Further cleavage causes the E-Syt3 relocation to the surface of LDs. Fragmentation of the primordial cisterna and LD budding into its lumen are early events in the biogenesis of LDs in the 3T3-L1 adipocyte. Electron tomography-based 3D reconstruction of the fragmented primordial cisterna revealed patches of a tightly packed E-Syt3-rich network of varicose tubules in close contact with young LDs. Esyt3 binds avidly phosphatidylethanolamine through its SMP domain, a main component of the LD membrane that fosters LD biogenesis. Repression of E-Syt3 effectively inhibits LD biogenesis and growth.

## Introduction

Lipid droplets (LDs) are dynamic cellular organelles that play essential roles in lipid metabolism, energy homeostasis, and lipid detoxification and trafficking as manufacturers and regulated reservoirs of triacylglycerols and sterol esters (SEs) (Martin and Parton, 2006; Walther and Farese, 2012). Recent investigations have implicated LDs in some unsuspected activities, including as platforms for lipid signaling in immunity and antibacterial response (Saka and Valdivia, 2012) and as repositories and lipidation sites for transcription factors and chromatin components (Haemmerle et al., 2011; Li et al., 2012).

LDs have a unique architecture consisting of a core of neutral lipids (NLs) formed by triacylglycerol/SE and wrapped in a phospholipid (PL) monolayer encapsulated by a protein coat. The PL monolayer originates from the endoplasmic reticulum (ER) membrane leaflet that wraps the budding nascent droplets, whereas the protein coat is assembled by selective protein-PL interactions mediated by a variety of hydrophobic mechanisms.

In non-adipose cells, the assembly of LDs has been shown to be confined to specific ER foci where metabolic enzymes, lipid substrates/products, and a variety of proteins that contribute to the assembly of LDs are gathered together (Chung et al., 2019; Kassan et al., 2013; Wang et al., 2016). How these foci are formed and whether they develop randomly or systematically through the ER is not known. In addition, few details are known about LD growth and the development of the giant LD that fills the cytoplasm in mature adipocytes.

The most accepted model of LD biogenesis, the lens model, proposes that the newly synthesized NLs coalesce between the ER membrane leaflets to create the lens-shaped seeds for new LDs (Murphy and Vance, 1999; Stein and Stein, 1967; Wanner et al., 1981). The lens model was indirectly supported by the capacity of bilayer membranes to hold NLs (King et al., 1994; Mackinnon et al., 1992), which tend to coalesce and form blisters (Khandelia et al., 2010), and by the continuity of the cytoplasmic leaflet of the ER membrane with the PL monolayer that wraps the LD (Ohsaki et al., 2017). Recently, the lens model was boosted by the unambiguous detection of lipid lenses within the ER membrane bilayer (Choudhary et al., 2015).

The limiting monolayer of the LDs is made by a complex mixture of PLs, the two most abundant of which are the glycerophospholipids (GPLs) phosphatidylethanolamine (PE) and phosphatidylcholine (PC). Changes in the PL composition of the ER membrane and LD monolayer dramatically affect LD biogenesis and growth. PE lowers membrane tension and fosters the rapid budding of small lipid droplets, whereas PC increases membrane tension and favors the slow formation of large LDs (Ben M’barek et al., 2017; Krahmer et al., 2011). In differentiating 3T3-L1 adipocytes, phosphatidylserine (PS) is imported from the ER and decarboxylated into PE in the mitochondria (MT); PE is then exported to the ER and LD. In the ER, PE fosters LD biogenesis and, on the LD surface, is converted into the PC required for LD growth and stability (Horl et al., 2011).

The separate sites of PE and PC production and action indicate a network of contacts between the MT, ER, and LDs. The contacts identified thus far include the membrane continuity between the ER and LD (Blanchette-Mackie et al., 1995; Jacquier et al., 2011; Salo et al., 2016) and the formation of transient junctions between the MT, ER, and LDs. Though the details of the assembly and regulation of the contacts are unclear, the use of a variety of proteins, including lipid transporters, in the complex task of establishing the contacts between LDs, the MT, and ER is becoming clear. Interestingly, ORP5, a protein localized to the ER/PM and ER/MT junctions and LDs, has been implicated in the transport of PS from the ER to the MT and in the control of PI4P levels in LDs (Du et al., 2020; Ghai et al., 2017; Renne and Emerling, 2020; Rochin et al., 2019). The role of ARF1/COP1 in the creation of LD/ER contacts (Wilfling et al., 2014) and SNAP23/PLIN5 in the formation of LD/MT contacts (Jagerstrom et al., 2009), as well as the use of seipin/LADF1 in the extraction of NLs that produce the LD foci in the ER (Chung et al., 2019), illustrate the variety of tools used in LD assembly.

The tricalbins in yeast and extended synaptotagmins (E-Syts) in mammalian cells function as tethers between the ER and plasma membrane (PM) (Giordano et al., 2013; Manford et al., 2012). E-Syts are ER membrane proteins with similar SMP and C2 domain organization as synaptotagmins. The E-Syt family has three members (Min et al., 2007): E-Syt1 has five C2 domains, and E-Syt2/3 each have three C2 domains. An N-hydrophobic hairpin anchors E-Syts to the outer ER membrane leaflet. E-Syt1 is associated with the inner ER, and an increase in cytosolic Ca^2+^ causes cortical relocation (Giordano et al., 2013). E-Syt2/3 are localized in the cortical ER and exploit positively charged patches in the C2C domain to bind PI(4,5)P_2_ and tether the ER to the PM. In vitro studies suggest that E-Syts exploit the dimerization of the SMP domain to exchange GPLs and diacylglycerol (DAG) between the apposed membranes (Saheki et al., 2016; Schauder et al., 2014).

The combined ability of E-Syts to tether the ER to the PM and to interact with activated FGF receptors has been proposed to increase FGF receptor endocytosis by clathrin-coated pits and FGF recycling (Herdman and Moss, 2016). In addition, loss of E-Syt2 and E-Syt3 affects in vitro cell migration and survival under stress (Herdman et al., 2014). Separately, E-Syt1 interacts with the glucose transporter GLUT4 upon phosphorylation by Cdk5 and may regulate its transport to the PM after insulin stimulation, as the suppression of Cdk5 activity inhibits glucose uptake by 3T3-L1 adipocytes (Lalioti et al., 2009).

Here, we report that E-Syt3 is truncated and relocates from the ER and ER-PM contact sites to a previously undescribed, single, giant annular structure, the primordial cisterna, when 3T3-L1 fibroblasts are induced to differentiate into adipocytes. Truncation is performed by proteasome-mediated proteolysis and results in removal of the C2C domain that links the full-length E-Syt3 to the PM. The primordial cisterna, an ER-derived structure consisting of a compact network of varicose tubules, mothers the LDs. E-Syt3 shows great affinity for GPLs (PE>PC) and participates in the biogenesis of LDs, process that is seriously hampered by E-Syt3 repression.

## Results

The differentiation of mouse 3T3-L1 fibroblasts into adipocytes is a widely validated model for studying the organelles and factors implicated in the biogenesis of LDs (Walther et al., 2017). E-Syt3 RNA is widely expressed in mouse non-adipose and adipose tissues as well as in 3T3-L1 adipocytes (Fig. S1). We used antibodies raised against E-Syt3 (Figs. S2, S3) and the ectopic expression of several E-Syt3 constructs to study the biogenesis of LDs.

### E-Syt3 localizes on the surface of LDs in 3T3-L1 adipocytes

In young adipocytes on day 3 of differentiation, E-Syt3 was associated with a population of small and medium-size LDs distinct from the small LDs coated by PLIN2 and PLIN3 (Fig. 1), proteins that are solely expressed in non-adipose cells and young adipocytes (Itabe et al., 2017). In grown adipocytes, the association of E-Syt3 with LDs overlapped with the population of droplets coated with PLIN1 (Fig. 1) (Brasaemle et al., 1997; Heid et al., 1998; Jiang and Serrero, 1992).

**Fig. 1.**
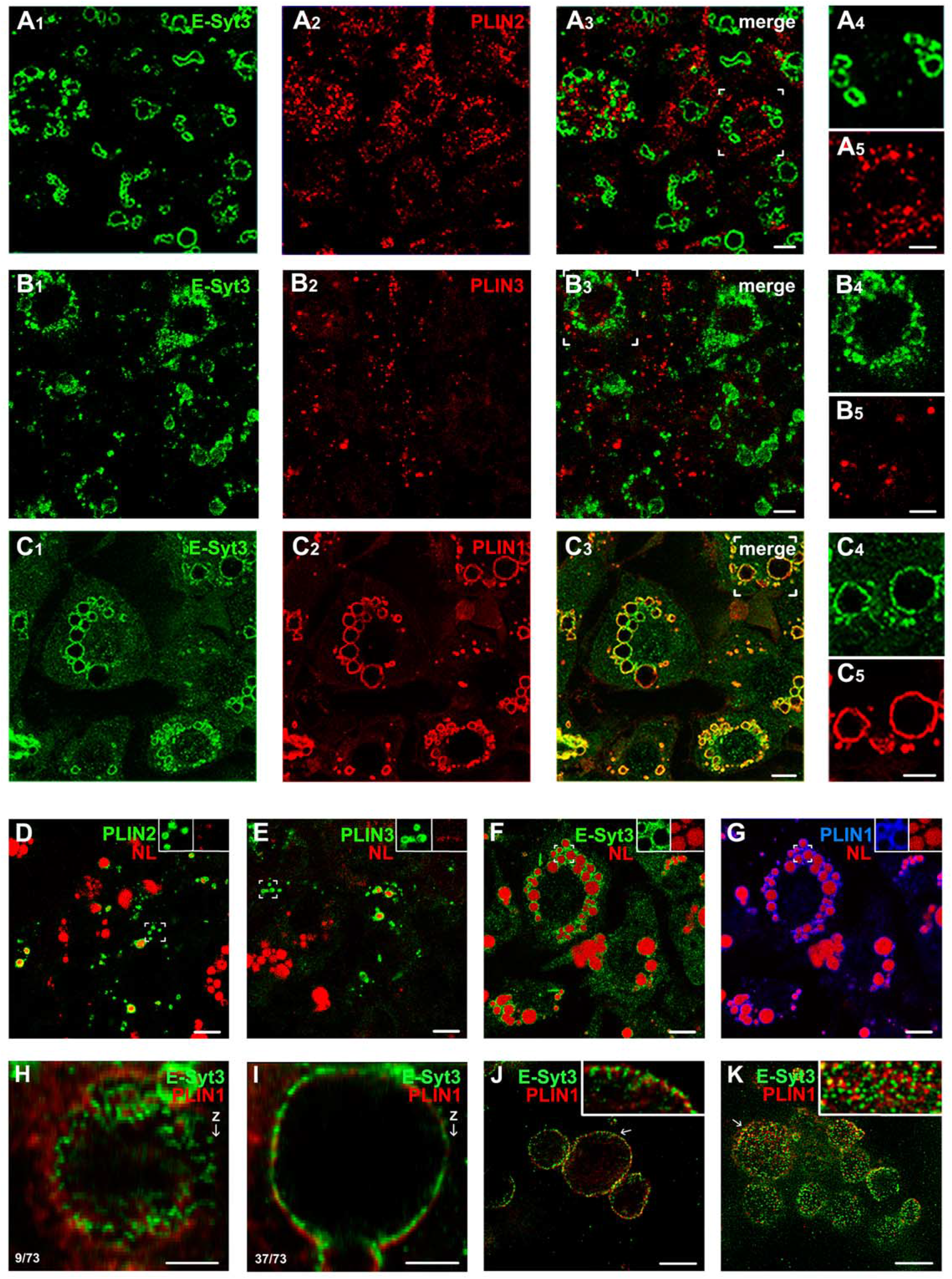
Subcellular distribution of endogenous E-Syt3 in 3T3 L1 adipocytes. Confocal images from 3T3 L1 adipocytes on days 3 (A, B, D, E) and 6 of differentiation (C, F, G). The localization of endogenous E-Syt3 was studied using affinity purified α-E-Syt3p^141^ antibody and specific PLIN1, PLIN2, and PLIN3 antibodies. The NL load of LDs was stained using LipidTOX. Note the absence of E-Syt3 in the PLIN2 (A_1_-A_5_) and PLIN3 (B_1_-B_5_) coated LDs, and its association with the PLIN1 positive LDs (C_1_-C_5_). Images D-G show the mutually exclusive staining of small PLIN2/PLIN3 droplets and medium/large PLIN1 droplets with the NL reagent LipidTOX. Areas framed by miter joints show enlarged magnifications of the LD images. High resolution confocal deconvoluted images (H, I) and STED images (J-K) showing the close association of E-Syt3 and PLIN1 with the LD surface and their different distribution, bottom left numbers indicate the section position within the stack. Scale bars = 6 μm (A-C), 10 μm (D-G), or 3 μm (H-K). The experiments were independently repeated at least three times on duplicate samples with the same results. Unless otherwise indicated, all endogenous E-Syt3/LDs microscopy were performed using the affinity purified rabbit/rat α-E-Syt3p^141^ antibodies.

### The C2C domain of E-Syt3 is removed by proteolytic processing sensitive to the proteasome inhibitor MG132

Previous studies in HeLa cells localized E-Syt3 in the cortical ER-PM junctions and showed that it is anchored to the PM by the interaction of the C2C domain with PI(4,5)P_2_, an especially abundant PIP2 in the PM (Giordano et al., 2013; Min et al., 2007). These studies also showed that removal of the C2C domain results in retention of the truncated protein in the inner ER. The association of endogenous E-Syt3 with LDs prompted us to examine whether the C2C domain was removed from the protein in adipocytes. To this end, we looked at the cleavage of C- and N-EGFP tagged E-Syt3 in transfected adipocytes (Figs. 2). Pilot experiments revealed significant C-cleavage of the two constructs in the fraction of heavy membranes and nuclei collected by low speed centrifugation (1000×*g*, 10 min). Therefore, we studied the cleavage in the nuclei-free membrane fraction extracted with 1% Triton X-100 and incubated for 5 min at 4°C. To prevent non-specific proteolysis, the incubation was performed in the presence of cOmplete Mini, a broad-spectrum inhibitor cocktail for lysosomes and metalloproteases.

**Fig. 2.**
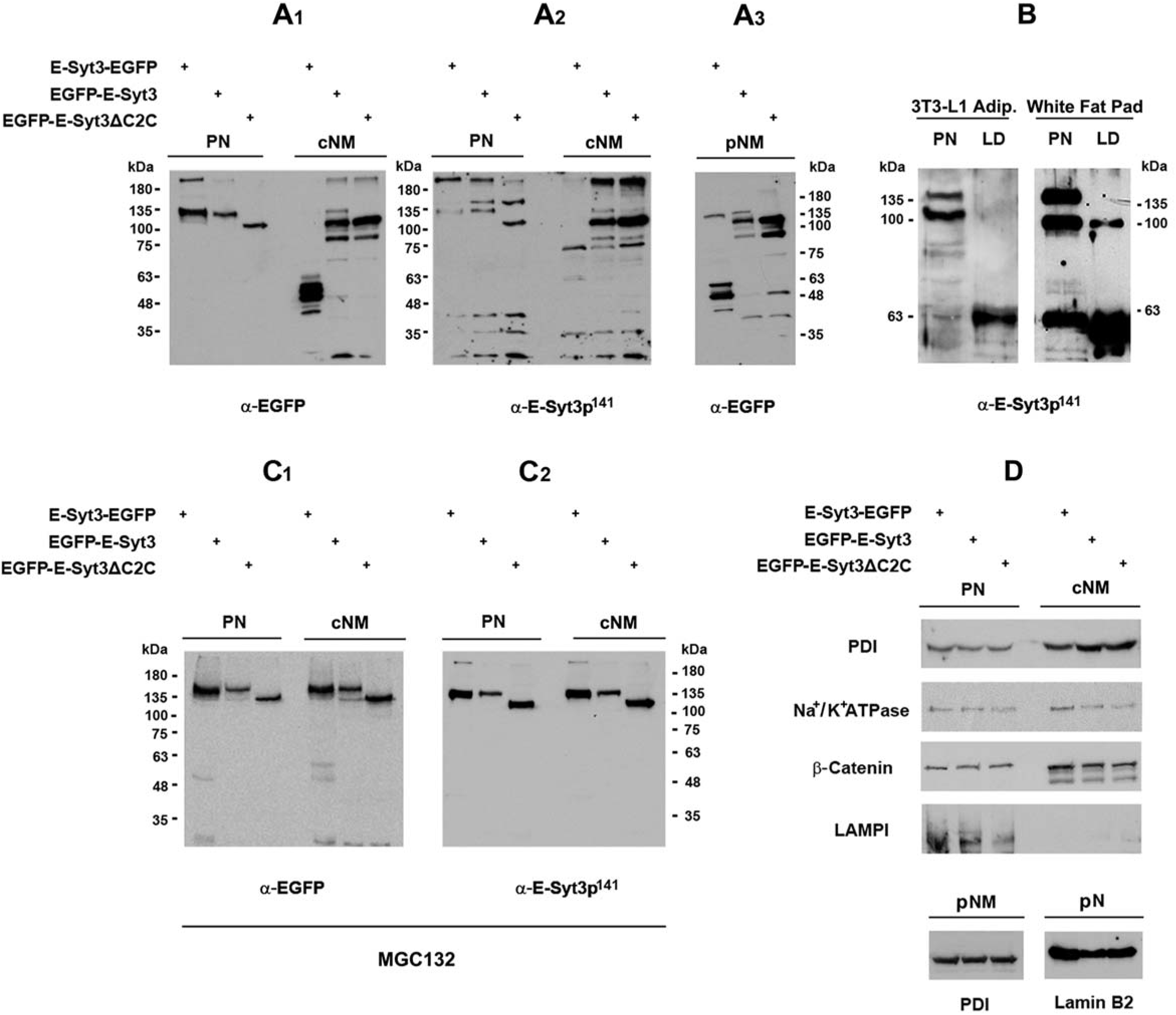
E-Syt3 cleavage and C2C domain removal. 3T3 L1 adipocytes were separately transfected for 36 h with E-Syt3 EGFP, EGFP E-Syt3, or EGFP E-Syt3ΔC2C to study the proteolytic processing of E-Syt3 by Western blotting. (A_1-3_) Cleavage of E-Syt3 in Triton X 100 extracts from postnuclear supernatants (PN, 20 μg) and crude (cNM, 20 μg) and purified (pNM 15 μg) heavy membranes collected by low speed centrifugation (1000×*g* 10 min). C- and N-E-Syt3 ends were identified using α EGFP and α-E-Syt3p^141^ antibodies. Note the absence of proteolytic processing in the PN fraction and the selective triple C-E-Syt3 cleavage in the cNM and pNM fractions. (B) Endogenous E-Syt3 species recovered in the PN fraction (10 μg) and purified LDs (15 μg) from 3T3 L1 adipocytes and white adipose tissue. Note the 100 kDa full-length E-Syt3 and 135 kDa species in the PN fraction and the 59‐63 kDa truncated E-Syt3 species in purified LDs. (C_1-2_) Inhibition of C E-Syt3 cleavage in transfected 3T3 L1 adipocytes treated the last 9 h of transfection with the proteasome inhibitor MG132 (50 μM). (D) Distribution of endoplasmic reticulum PDI, plasma membrane Na^+^/K^+^ ATPase, cytoplasmic/nuclear β-catenin, lysosomal LAMP1, and nuclear membrane lamin B2 among the PN, cNM, pNM, and pN (purified nuclei) fractions. The studies were repeated three times with similar results.

Western blot analysis of the proteolytic digestion of E-Syt3-EGFP using an anti-EGFP antibody showed that the protein was neatly cleaved into three major C-peptides (60, 55, and 50 kDa) carrying the C2C domain within E-Syt3 fragments of 32, 27, and 22 kDa (Fig. 2A_1_). Study of the resulting complementary N-fragments in extracts from cells transfected with EGFP-E-Syt3 resulted in identification of two major peptides of 106 kDa and 87 kDa, and a minor peptide of 101 kDa, that carried E-Syt3 fragments of 78, 59, and 73 kDa, respectively (Fig. 2A_1-3_). The 78/22 kDa and 73/27 kDa pairs accounted for the 100 kDa full-length E-Syt3, whereas the sum of the 59/32 kDa pair produced a 91 kDa peptide, 9 kDa shorter than expected. The recovery of the 59 kDa peptide strongly suggests that the expected 68 kDa species was cleaved into peptides of 59 kDa and 9 kDa (see Fig.S4 for a complete description of the stepped protein cleavage). In an attempt to study in more detail the cleavage, we artificially produced a 106 kDa truncated EGFP-E-Syt3ΔC2C protein (Fig. S1). Both, the transfected EGFP-E-Syt3ΔC2C and the co-migrating 106 kDa EGFP-E-Syt3 proteolytic product were cleaved into an 87 kDa peptide (Fig. 2A_1-3_, Fig. S4), indicating that the natural and artificial cleavage sites mapped closely, proximity that validated the experimental use of EGFP-E-Syt3ΔC2C.

The recovery of 59 E-Syt3 species with highly purified LDs from 3T3-L1 adipocytes and white fat adipose tissue (Fig. 2B) established a cause-effect relationship between the C-cleavage and the association of E-Syt3 with LDs in adipocytes. Furthermore, we observed that incubation of 3T3-L1 adipocytes with the proteasome inhibitor MG132 strongly inhibited the stepped C-cleavage of the EGFP-tagged E-Syt3 constructs (Fig. 2C_1-2_). Moreover, study of the E-Syt3 ubiquitination in immunoprecipitates using α-E-Syt3p141 and anti-ubiquitin antibodies, showed that whereas truncated E-Syt3 was hardly ubiquitinated, two E-Syt3 species of 90 kDa and 180 kDa produced a clear ubiquitinated signal (Fig. S5). With regard to this it is noteworthy that a 180 kDa E-Syt3 species was recovered with purified LDs (Fig. 2C). Taken together, these results point to a role of the ubiquitin/proteasome system in the proteolytic processing of E-Syt3.

### Truncation of the C2C domain results in LD targeting by EGFP-tagged E-Syt3 in 3T3-L1 adipocytes

Next, we compared the cellular distribution of EGFP-E-Syt3 and EGFP-E-Syt3ΔC2C (Fig. 3A) in 3T3-L1 fibroblasts and adipocytes. In agreement with the previous study by Giordano et al. (2013), in 3T3-L1 fibroblasts transfected with EGFP-E-Syt3, the protein was retained in the ER and in punctate structures on the cell surface, the typical pattern of PM/ER junctions (Fig. 3B_1-2_). As expected, truncated EGFP-E-Syt3ΔC2C was completely retained in the ER (Figs. 3C, S6A).

**Fig. 3.**
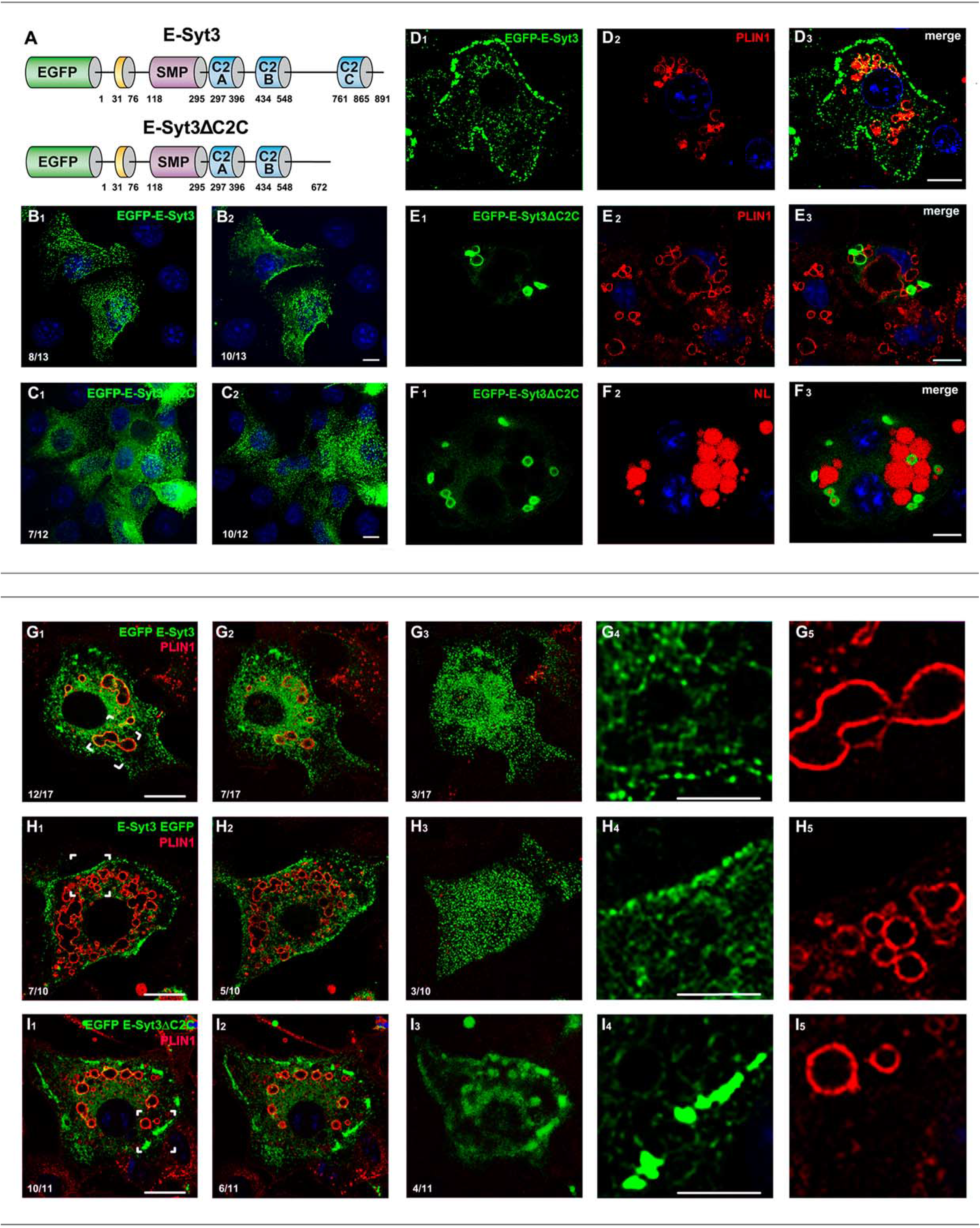
Subcellular distribution of EGFP tagged E-Syt3 and E-Syt3ΔC2C in transfected 3T3-L1 fibroblasts and 3T3-L1 adipocytes. (A) Schematic showing the domain structure of the full length and C truncated EGFP tagged E-Syt3 constructs. The N hydrophobic hairpin that anchors the protein to the ER membrane (yellow) is sequentially followed by the SMP domain involved in the transport of lipids and three C2 domains. EGFP-E-Syt3ΔC2C was produced by removal of the C2C domain. (B-C) The cellular distribution of transfected EGFP-E-Syt3 and EGFP E-Syt3ΔC2C in 3T3 L1 fibroblasts is evident in 0.8 μm optical sections taken through the cell interior (B_1_, C_1_) and the plane of cell attachment to the substrate (B_2_, C_2_). Note the contrast between the association of EGFP E-Syt3 with the cell surface and the absence of EGFP-E-Syt3ΔC2C (see also Fig. S3A). (D-F) Cellular distribution of EGFP-E-Syt3 and EGFP-E-Syt3ΔC2C in adipocytes on day 5 of differentiation. Note the association of EGFP-E-Syt3 with the PM and its partial localization in LDs (D1 3). In contrast, observe the association of EGFP-E-Syt3ΔC2C with PLIN1 (E_1-3_) and NL rich LDs (F_1-3_), and with a few oblong PLIN1/NL negative structures (see also Fig. 6A). NLs were stained with LipidTOX and nuclei with DAPI. The experiments were independently repeated at least three times on duplicate samples with comparable results. Scale bars = 10 μm. (G-I) Proteasome inhibition impedes the incorporation of transfected EGFP E-Syt3 and EGFP E-Syt3ΔC2C into LDs. In the subcellular distribution of full length and C truncated EGFP tagged E-Syt3 constructs in 3T3 L1 adipocytes treated with the proteasome inhibitor MG132 (50 μM, 9 h), note the restriction of EGFP E-Syt3 (G_1-5_) and E-Syt3 EGFP (H_1-5_) localization to the ER and the punctate ER/PM junctions and the retention of E-Syt3ΔC2C within a cortical segmented cisterna like structure (I_1-5_). The areas framed by miter joints are enlarged in panels G_4-5_, H_4-5_, and I_4-5_. Compare the images shown in panels I_1-5_ with the live cell time lapse fluorescence microscopy images recorded in videos 1 and 2. Scale bars = 20 μm (5 μm in enlargements). Bottom left numbers indicate the section position within the stack. All of the experiments were independently repeated three times on duplicate samples with similar results.

In 3T3-L1 adipocytes, EGFP-E-Syt3 was mainly associated with ER/PM junctions and a small fraction with LDs (Fig. 3D). In sharp contrast, E-Syt3ΔC2C exclusively targeted a fraction of the small/medium-sized PLIN1/NL-positive LDs and a few oblong PLIN1-negative cisterna-like structures with few or no NLs (Fig. 3E, F; see below Fig. 6A, B). To note, removal of the 35 amino acid neosequence that extends the C-end of EGFP-E-Syt3ΔC2C (Fig. S1) did not change its distribution (Figs. S6B, C). The exclusive localization of E-Syt3ΔC2C in LDs and cisternae indicated that the C-truncation simultaneously blocked the targeting of the protein to the cell surface. Remarkably, the truncated protein was not found in large LDs, strongly suggesting that it was not incorporated into pre-existing LDs. These results and the recovery of the 63 kDa E-Syt3 species in purified LDs (Fig. 2B) indicate that the targeting of endogenous E-Syt3 to LDs is preceded by cleavage of the C2C domain.

Due to the sequence homology between the segments containing the SMP-C2A-C2B domains of E-Syt3 and E-Syt1 and their regulated cooperation in the ER/PM junctions in non-adipose tissue, we studied the possibility that E-Syt1 is C-cleaved and targeted to LDs using an α-E-Syt1 antibody (Figs. S7). Unlike EGFP-E-Syt3ΔC2C, EGFP-E-Syt1ΔC2C-E was retained in the ER (Fig. S6D), suggesting that specific motifs retain truncated E-Syt1 in the ER or target EGFP-E-Syt3ΔC2C to LDs.

As inhibition of the proteasome completely blocks C-cleavage of the full length and the ΔC2C protein, we investigated whether this changed their subcellular distribution. MG132 treatment blocked their targeting of LDs. Moreover, whereas the retention of full-length protein in the ER and in the ER/PM junctions was not affected (Fig. 3G, H), the MG132 treatment caused retention of the majority of EGFP-E-Syt3ΔC2C in a discontinuous cisterna-like structure localized in the cell cortex (Fig. 3I). This observation suggested that the cortical cisterna was in the path of the protein to LDs, and that further proteolytic processing of the truncated protein was necessary to reach the LDs. The link between the E-Syt3 cleavage and its cellular localization led us to study the early steps of LD formation by live-cell time-lapse IFM microcopy, using the fluorescent truncated protein as a tracer.

### E-Syt3 localizes to a single, large, circular cisterna before translocating to the LD surface

In eukaryotic non-adipose cells, LDs are synthesized in restricted foci/regions in the ER. In the foci are recruited the lipid synthesis enzymes and assembly factors involved in the formation of the NL lenses that modify the membrane from which nascent LDs bud (Henne et al., 2019; Jacquier et al., 2013; Kassan et al., 2013; Wang et al., 2016; Wang et al., 2018). As shown in live-cell time-lapse IMF microscopy, in 3T3-L1 adipocytes the targeting of EGFP-E-Syt3ΔC2C to the surface of newly formed LDs was preceded by its transient appearance in a discontinuous cortical cisterna similar to the structure observed in adipocytes treated with MG132 (Videos 1, 00.40-01.30 h; video 2, 11.30-11.50 h; Fig. 3I). To investigate the cisterna in untreated cells, we monitored changes in the distribution of endogenous E-Syt3 in fixed young adipocytes on days 1-3 of differentiation using confocal IFM. Although this approach was challenged by the relaxed synchrony of the differentiation process and the 10-20 min window available to study the cisterna (Videos 1, 2), we expected that, if successful, it could provide the good-quality images necessary to confirm the cisterna existence and to study this in the context of LD biogenesis. Within a field crowded with cells containing collars of small and medium-size LDs, we found a small group of cells with endogenous E-Syt3 retained in a long, circular, cortical cisterna. The cisterna, which has an average perimeter of 74 ± 9.5 μm, was either branched or smooth, and was embedded in the vast ER (Fig. 4A-B, Fig. S8A-C).

**Fig. 4.**
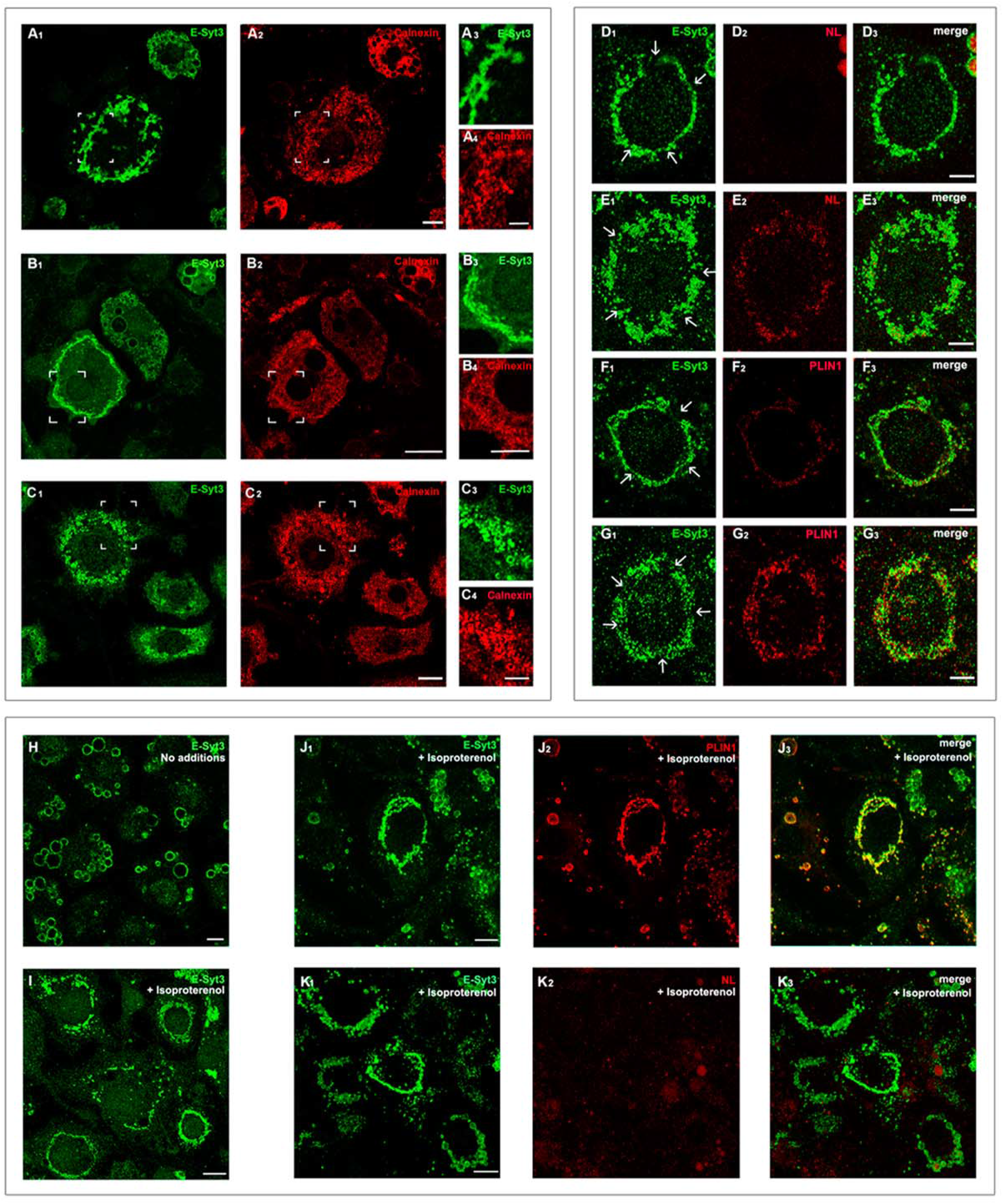
Relocation of endogenous E-Syt3 during early adipocyte differentiation. (A-C) Confocal microscopy of preadipocytes on day 3 of differentiation. Cells were fixed, permeabilized, and stained using α E-Syt3p^141^ and α calnexin antibodies. Note the E-Syt3 positive circular/branched cisternae embedded in the calnexin positive ER (A_1-4_), the smothering of the cisterna (B_1-4_), and its subsequent fragmentation into clusters of polymorphic cisternal structures (C_1-4_). The miter joints demarcate the enlarged areas (A_3-4_, B_3-4_, C_3-4_). Scales bars = 20 μm (10 μm in enlargements). Observe that the recruitment of E-Syt3 and PLIN1 into the primordial cisterna precedes NL synthesis (compare D_1_, F_1_ and F_2_ with D_2_) and that fragmentation of the cisterna coincides with the formation of the first E-Syt3/PLIN1/NL positive LDs. Scale bars = 9 μm. Lipolysis causes the replacement of E-Syt3 positive LDs by a NL free primordial cisterna like structure. On day 6 of differentiation, adipocytes were untreated (H) or starved and treated overnight with 10 μM isoproterenol (I-K). Note the the substitution of LDs by collars of E-Syt3/PLIN1 positive small droplets and cisternae (I, J_1-3_) with negligible levels of NLs (K_1-3_). Scale bars = 10 μm. The experiments were independently repeated three times on duplicate samples with comparable results.

With regard to the relocation of EGFP-E-Syt3ΔC2C from the cisterna to LDs, an event that is tightly connected to LD biogenesis, we observed initial colocalization with the ER membrane protein calnexin in the cisterna and newborn LDs (Figs. 4B-C) and an absence of calnexin from the medium/large LDs (Fig. S9B). This transient colocalization and embedding of the cisterna in the ER are both consistent with the primordial cisterna being a medium-size region (10-20%) of the ER specialized in LD biogenesis. Of particular interest was the conversion of the continuous cisterna (Fig. 4A, B) into the segmented structure observed in transfected adipocytes treated with MG132 (Fig. 3I) and in live-cell time-lapse IFM experiments (Videos 1 and 2, Fig. 3I). The segmentation does not appears to result from the unequal distribution of the protein within the cisterna, since it was followed by an extensive structural reorganization, including further fragmentation that coincided with the appearance of the first NLs drops (Fig. 4C-, E, G). Importantly, E-Syt3 and PLIN1 were recruited into the cisterna before the detection of the first NL drops (Fig. 4D-G). This observation is consistent with the cisterna being the site where the materials and machinery necessary for the assembly of the LDs are recruited.

The adipocyte is subjected to repeated cycles of NL synthesis and degradation in the context of the lipid/carbohydrate homeostasis. We used E-Syt3 and PLIN1 as tracers to study if after lipolysis the collapse of the LDs was followed by the reappearance of the primordial cisterna. On day 6 of differentiation, adipocytes were starved and incubated overnight with 10 μM isoproterenol. Strikingly, the digestion of the NLs resulted in substitution of the large and seemingly individual LDs by a continuous E-Syt3/PLIN1-positive, NL-negative, cisterna similar to the original primordial cisterna (Fig. 4I-K). This substitution might be the result of the recycling of materials through the primordial cisterna, for their use in the synthesis of new LDs, or indicate that LDs are expansions of the primordial cisterna that collapse into this after lipolysis, alternatives that are not mutually exclusive (see Discussion).

### Esyt3 is associated with patches of a tightly packed network of varicose tubules associated with young LDs

To gain further insight into the ultrastructural details of the primordial cisterna, we studied young adipocytes transfected with EGFP-E-Syt3ΔC2C using correlative light/immunoelectron microscopy (Figs. 5, S10). We examined cells in the early phase immediately before concentration of the truncated protein in the cisterna (Fig. S10, stage 1), and upon its concentration into the globular cisterna segments from which the LDs grow (Fig. S10, stage 4). Before its concentration in the cisterna, the protein was associated with smooth ER elements circularly arranged in the cytoplasm (Fig. 5A). Later, and coinciding with the formation of the first LDs, the protein was concentrated in contiguous patches of tightly packed smooth ER tubules, wrapped by a membrane, in the proximity of clustered young LDs (Fig. 5B). Interestingly, 3D electron tomography reconstruction of these tubular patches revealed a highly dense network of varicose tubules whose membranes were regularly beaded (Fig. 5C_1-3_) (see Discussion).

**Fig. 5.**
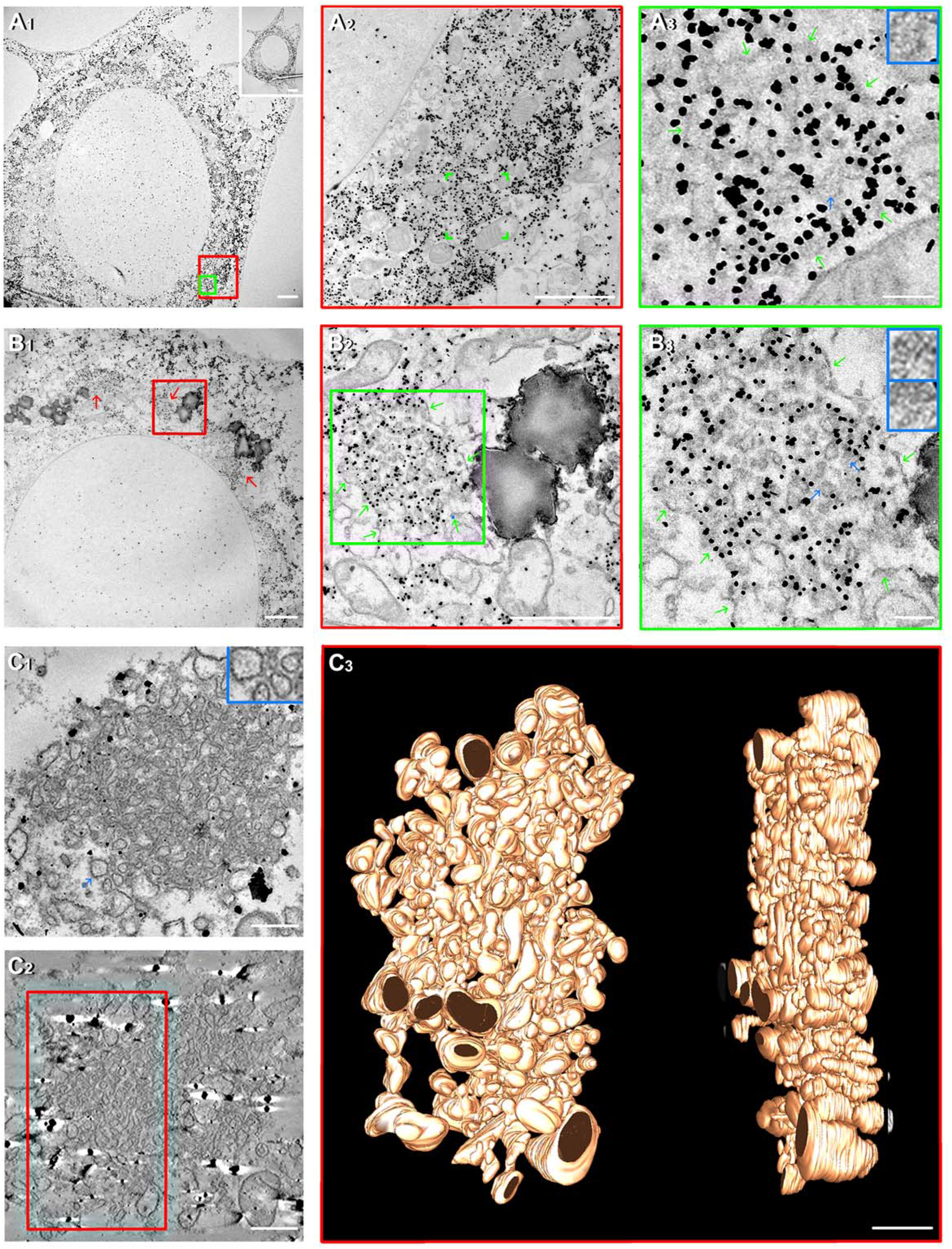
Electron microscopy study of the EGFP-E-Syt3ΔC2C distribution in preadipocytes and young adipocytes: Electron tomography (ET) based 3D reconstruction of cisterna segments. (A) Young adipocytes at stage 1, a phase that precedes the accumulation of EGFP-E-Syt3ΔC2C in the primordial cisternae, and (B,C) stage 4, upon the cisterna segmentation that coincides with the appearance of the first LDs, were selected as described in Fig. S10 (white arrows) and the subcellular areas (red arrows) studied by high resolution EM using a rabbit anti EGFP antibody. Note the annular arrangement of gold labelled EGFP E-Syt3ΔC2C in the young adipocyte at stage 1 (A_1_, A_2_) and its association with tubular ER elements (A_3_) with vesicle like protrusions (14.4 ±2 nm; blue arrow, insert; green arrow, wrapping membrane). In young adipocytes at stage 4 (B, C), note the retention of gold labelled EGFP-E-Syt3ΔC2C within foci consisting of contiguous/globular cisterna segments (48.5±12 μm) (B_1_, red arrows) associated with small clusters of young LDs (0.3 μm) (B_1, 2_; green arrow, cluster membrane). In addition, note the filling of the cisterna segments with densely packed tubules (B_2_, B_3_, compare with A_3_) and their beaded membranes (blue arrows, inserts). High resolution EM of two 70 nm (C_1_) and 200 nm (C_2_) consecutive sections of the cisterna segment (red arrow) in Fig S10C_1_. Note the beaded tubular membranes (C_1_, insert). ET study of the 200 nm section (C_3_); observe the densely packed network of varicose tubules in the 3D reconstruction. Scale bars = 2 μm (A_1_, B_1_, C_3_), 1 μm (A_2_, B_2_), and 200 nm (A_3_, B_3_, C_1_, C_2_).

### E-Syt3 association with LDs precursors and growing LDs

On day 4 of differentiation, monitoring the distribution of EGFP-E-Syt3ΔC2C in 3T3-L1 adipocytes by simultaneous immunofluorescence and bright field microscopy using live-cell time-lapse fluorescence microscopy showed the appearance of a diffuse fluorescence. This fluorescence was concentrated into several cisternae, and then into one cisterna, before its apparent fragmentation into several oblong structures loaded with small LDs (Videos 3, 4). Comparing the IFM and bright-field microscopy recordings revealed encapsulation of 1, 2, or 3 LDs (0.11 ± 0.02 μm) by a birefringent material containing the fluorescent EGFP-E-Syt3ΔC2C (Fig. 6A). Moreover, the encapsulated LDs fused with each other to gain body within the lumen of the 0.36 ± 0.12 μm diameter encircling membrane structures (Fig. 6C), structures that also fused with each other (Fig. 6B). These observations were unexpected and strongly suggest that, in 3T3-L1 adipocytes, the inward budding of LDs into the lumen of the ER plays an unsuspected role in LD biogenesis (Mishra et al., 2016).

**Fig. 6.**
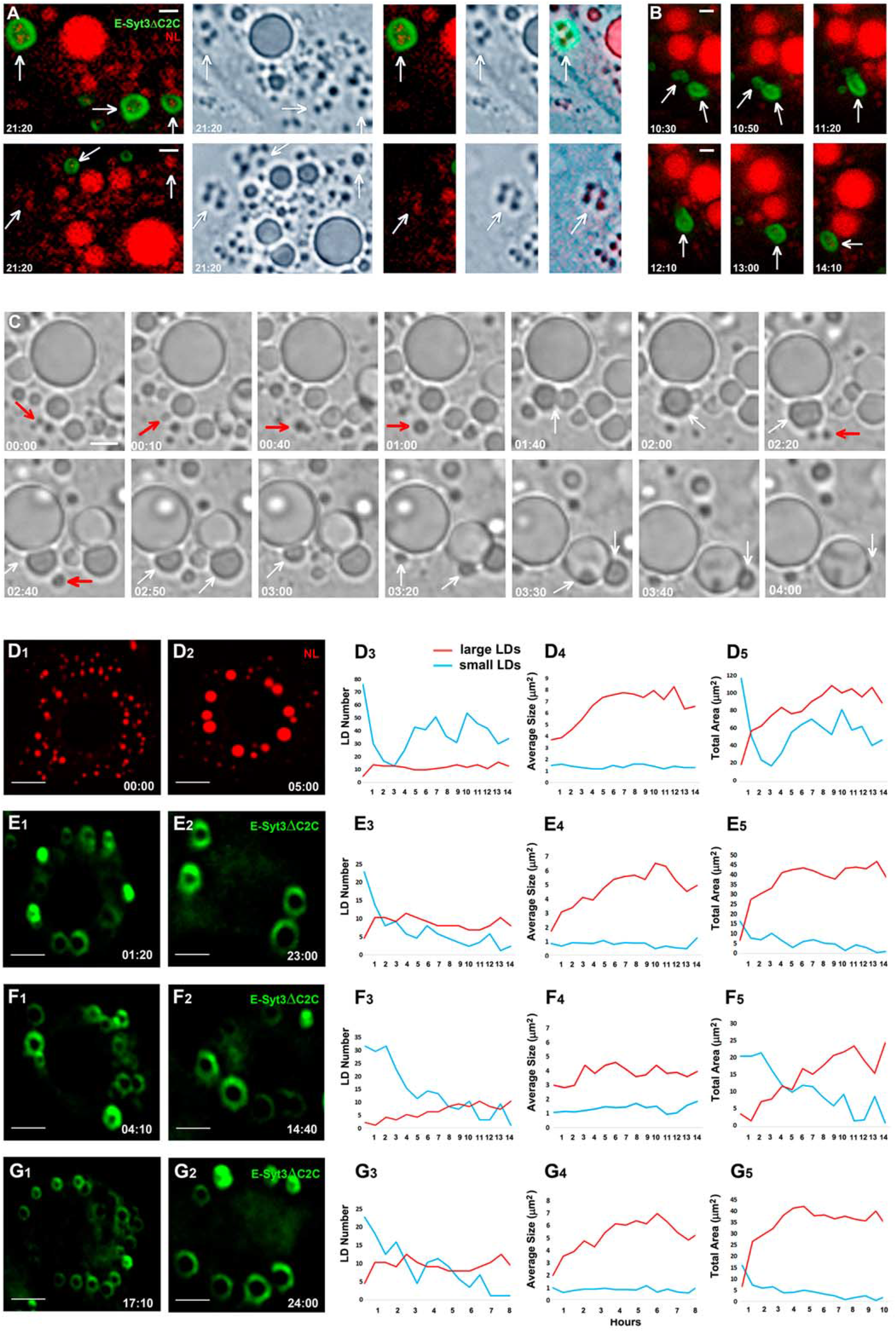
E-Syt3ΔC2C redistribution in transfected adipocytes through the early steps of LD assembly and growth. Growth of E-Syt3ΔC2C coated lipid droplets by homotypic fusion. Live cell time lapse fluorescence/bright field microscopy of adipocytes transfected with EGFP-E-Syt3ΔC2C on day 4 of differentiation. Selected paired (A, B) and single (C) images from videos 3 and 4 show the encapsulation of newborn LDs within cisterna like structures containing the truncated fluorescent protein laid over the birefringent capsule (A, arrows); due to the loss of small amounts of NLs during the cell permeabilization/fixation it is uncertain to asses if the absence of NL in some cisternae was real (see Fig 3F). Note the ongoing fusion between the cisternae (B, arrows), as well as between the encapsulated LDs (C, red arrows), and the predation of small LDs by the large LDs (C, white arrows). Scale bars = 2.65 μm (A), 1.97 μm (B), or 3.76 μm (C). (D-G) The dynamics of LD fusion in control adipocytes stained with LipidTOX (D, video 6) and in adipocytes transfected with EGFP E-Syt3ΔC2C (E-G, video 5). Snapshots show the LDs at the indicated recording times (right bottom). Line graphs represent the fusion profiles. Observe the fluorescent rings produced by the incorporation of EGFP E-Syt3ΔC2C into the surface of the LDs (E_1-2_; F_1-2_; G_1-2_; Video 6). Fusion events were quantified by measuring the changes in the number, average size, and total area in two separate populations of small (<3 μm) and large (> 3 μm) LDs (D_3-5_ G_3-5_). Note the initial prevalence of the small LDs and their rapid decrease (blue) as a result of the predatory activity of the large LDs (red). See the gradual slowdown in the fusion activity of the large LDs after reaching the 5-8 μm size and the variable production of new waves of small LDs. The experiments were repeated at least three times. Scale bars = 7.53 μm.

Concerning the localization of EGFP-E-Syt3ΔC2C in maturing adipocytes, the production of large fluorescent rings (Video 5, Fig. 6E_1-2_, F_1-2_, G_1-2_) agreed with the localization of endogenous E-Syt3 on the surface of LDs (Fig. 1). The LDs coated with the truncated protein exhibited the same dynamism and tendency to fuse with each other as their peers in control adipocytes stained with LipidTOX (compare Videos 5 and 6) indicating, therefore, that truncated E-Syt3 is tightly associated with the surface of mobile LDs. This conclusion, which excludes the retention of truncated E-Syt3 in the ER that encircles the LDs, is also supported by the distinct subcellular distribution of EGFP-E-Syt3ΔC2C and five different ER membrane proteins (Fig. S9) and the contrast with the relative motionless behavior of ER membrane marker EGFP-Sec61β (Video 7).

Monitoring LD growth revealed a process of continuous homotypic fusion that included the fusion of small LDs with each other and their subsequent predation by the large LDs (Fig. 6C-G; Videos 5, 6). Typically, under our experimental conditions, the 5-8 μm LDs stopped growing (Fig. 6D-G). The high number of fusion events recorded in our studies indicated the important contribution of homotypic fusion to LD growth.

The grown LDs were arranged in collars that retained the circular profile of the primordial cisterna and nascent LDs suggesting that the cisterna might fix the position of LDs (Fig. S8D, E) (see Discussion). Maintaining the adipocytes for 3 weeks in culture resulted in the formation of single-giant LDs that resembled the LDs found in adipose tissue (Fig. S8F, G).

### Phosphatidylethanolamine and phosphatidylcholine strongly bind to the E-Syt3 SMP domain

The detection of GPLs and DAG in the channel made by the purported dimerization of the E-Syt2 SMP domain suggests that E-Syts play an important role in the exchange of these materials between the apposed ER and PM lipid bilayers (Schauder et al., 2014). This experimentally based hypothesis, together with the critical roles played by PE in the control of LD biogenesis and PC in LD growth (Ben M’barek et al., 2017; Horl et al., 2011), made interesting to study the binding of PLs to the SMP and C2 domains of E-Syt3. Among the 16 PLs tested, PE and PC were the only ones to bind the SMP domain (Fig. 7A). The binding was strong and specific, and the binding of PE prevailed over PC binding. In contrast, phosphatidylinositol monophosphates and diphosphates (PPIns) solely and weakly bound to the C2B and C2C domains.

**Fig. 7.**
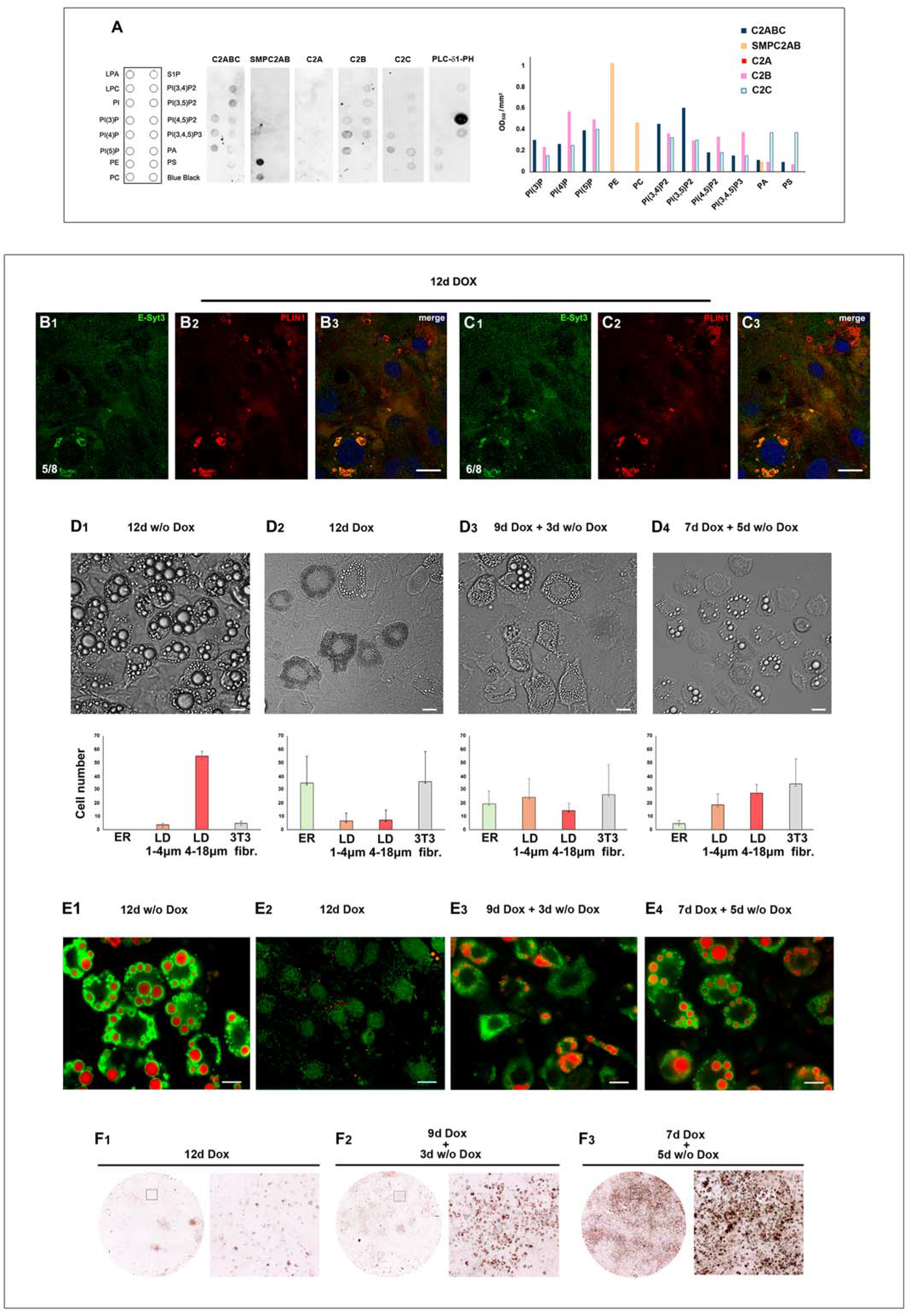
Repression of E-Syt3 expression by shRNA interferes with LD biogenesis. (A) The E-Syt3 SMP domain binds selectively and strongly PE and to a less extent PC. PIP Strip membranes were incubated with the GST conjugated/soluble C2ABC, SMPC2AB, C2A, C2B, and C2C domains from E-Syt3 (1 μg) and control PLC-δ1-PH. Note the selective and strong binding of PE>PC to the SMP domain and the relative weak reaction of PPIns with the C2B and C2C domains. The PIP experiment was repeated twice with comparable results. For E-Syt3 shRNA interference, non confluent 3T3-L1 fibroblasts were infected with E-Syt3 shRNA/lentivirus. The transduced cells were cloned (B, C) and subjected to adipocyte differentiation in the absence or presence of 200 ng/ml doxycycline (Dox) for 12 days to repress the expression of E-Syt3. Cells treated for 9 or 7 days with Dox were incubated for 3 or 5 days without Dox to reverse the repression. At the end of the indicated incubation period the expression and subcellular localization of E-Syt3 was studied by confocal (B, C) and epifluorescence microscopy (E), and the cell morphology and number and size of the LDs was quantified using bright field microscopy (D). NLs were stained with LipidTOX (E) or using the Oil Red O Lipid Stain kit (F). In the cells incubated for 12 days with Dox the formation of a broad ER collar is in stark contrast to the cell background (D_2_), note the strong reduction in E-Syt3 expression and inhibition of LD biogenesis (B-C, E_2_). Observe the partial recovery of LD biogenesis and renewed expression of E-Syt3 (D_3-4_, E_3**-4**_) and NL synthesis (E_3-4_, F_2-3_) following the removal of Dox. The experiments were repeated three times on duplicate samples with comparable results. Scale bars = 10 μm. B, C, bottom left numbers indicate the section position within the stack.

### Silencing of E-Syt3 strongly inhibits LD biogenesis

Next, we studied whether E-Syt3 repression would alter LD biogenesis. An inhibitory effect was clear; in clonal cells subjected to E-Syt3 shRNA interference, the formation of LDs was almost suppressed and adipocyte differentiation was poor. Though a few cells assembled a small number of medium-sized PLIN1/E-Syt3-positive LDs, the majority of the residually expressed E-Syt3 did not co-distribute with the PLIN1 preferentially associated with small LDs (Fig. 7B-C). Remarkably, the E-Syt3-negative cells observed by bright-field microscopy often exhibited the typical dense-broad ER collars from which LDs bud (compare Figs. 7D_2_ and 4C-G), an indication that the E-Syt3 repression interfered with the early steps of LD biogenesis. The inhibition was partly reversible (Fig. 7D, E) and resulted in the recovery of NL synthesis (Fig. 7E, F) and the resumption of LD biogenesis, resumption that was initiated with the accumulation of large amounts of small LDs in the cytoplasm, followed by their growth (Fig. 7D_3-4_).

## Discussion

Earlier studies investigating E-Syts in mammalian cells have mainly focused on their role in tethering the ER to the PM and in the exchange of lipids between the two apposed membranes. Consequently, it is not known whether E-Syts are also involved in the contact and exchange of lipids between the ER and other organelles and their physiological significance.

In a previous study of the distribution of E-Syt3 in cells from non-adipose tissues, artificial removal of the C2C domain resulted in intracellular retention of the protein within the inner ER and in detachment of the ER anchored to the PM (Giordano et al., 2013). In 3T3-L1 adipocytes, we found that the C2C domain is removed by proteolytic cleavage controlled by the proteasome, and that the resulting proteolytic product is recovered with highly purified LDs. These findings explain the novel association of endogenous E-Syt3 with LDs in 3T3-L1 adipocytes.

We monitored the proteolytic processing and subcellular distribution of full-length E-Syt3 and artificially truncated E-Syt3ΔC2C during adipocyte differentiation to study LD biogenesis and growth. The retention of full-length E-Syt3 in the ER and ER/PM junctions and EGFP-E-Syt3ΔC2C in a cisterna-like cortical structure in adipocytes treated with the proteasome inhibitor MG132 led us to identifying the primordial cisterna, an organelle with origins in the ER involved in LD biogenesis. This different localization strongly suggest that E-Syt3 is cleaved first to facilitate its access to the primordial cisterna and then to enable targeting from the cisterna to LDs. This agrees with the recovery of a truncated 59-63 kDa species with purified LDs. Furthermore, the precised step cleavage of the protein and its resistance to the cOmplete Mini anti-protease cocktail are indications of the specificity of the proteolytic cleavage. Further studies are warranted to establish whether E-Syt3 is cleaved by the proteasome itself or by proteases under its control as well as their regulation.

The targeting of E-Syt3ΔC2C to the LD surface is in contrast with retention of the C-truncated E-Syt1ΔC2C, D, E in the ER. Both E-Syt3 and E-Syt1 have N-hydrophobic hairpins that are used for anchoring to the cytoplasmic leaflet of the ER anchor. Hydrophobic hairpins are used by some coat proteins to traffic and hang on the LD surface (Ohsaki et al., 2017; Wilfling et al., 2013). Therefore, the difference between the location of E-Syt1 and E-Syt3 in 3T3-L1 adipocytes suggests the existence of additional motifs that retain E-Syt1 in the ER or target E-Syt3 to LDs.

Studies in cells from non-adipose tissues have shown that LDs are assembled in specific microdomains of the ER called foci (Kassan et al., 2013; Wang et al., 2016). The finding that LDs are generated from a single giant cisterna in 3T3-L1 adipocytes was unexpected. The live-cell time-lapse studies of adipocytes transfected with EGFP-E-Syt3ΔC2C showed that the intact primordial cisterna is briefly visible for a period of 10-20 min. This short time window and the asynchrony of the process of adipocyte differentiation explain why the cisterna is much less frequently seen than its offspring LDs in preparations of fixed cells and may explain why it was not detected before.

The intact primordial cisterna is E-Syt3/PLIN1/calnexin positive, but it did not stain with LipidTOX. Though we do not know the details of the relationship between the cisterna and the vast and multifunctional ER, the presence of calnexin in the cisterna indicates its ER origin. Furthermore, the accumulation of E-Syt3 and PLIN1 in the primordial cisterna before the appearance of the first LDs strongly suggests that the cisterna is the organelle where the proteins that participate in LD formation are reunited. It is logical to expect that the primordial cisterna is produced in the ER via molecular aggregation initiated with the synthesis of necessary nucleation factors and propagated with the recruitment of all materials necessary for LD formation.

With regard to the dynamics of the primordial cisterna, live-cell microscopy studies using EGFPΔC2C showed that the fluorescent rapidly becomes segmented. The independent movement of the fluorescent segments and changing proximity to one another, as well as our failure to find any evidence that they are linked together, suggest that the cisterna is physically fragmented. The cisterna breaking may result in disconnection from the rest of the ER avoiding any undesirable interference between the two parts.

Staining of the fragmented cisterna with LipidTOX revealed a mixture of NL-positive and negative fragments, consistent with the asynchronous beginning of LD biogenesis.The appearance of nascent LDs in the lumen of the cisterna fragments, followed by their fusion into small LDs and the subsequent predation of these by large LDs, appear to describe the process of LD growth in 3T3-L1 adipocytes. Probing the topology of LD biogenesis by targeting LD-resident proteins to the ER lumen results in recovery of these proteins in cytoplasmic LDs, suggesting an ER domain that is accessible from the cytosolic and ER luminal side and the possible budding of LD into the ER lumen before appearance in the cytoplasm (Mishra et al., 2016). Clearly, further studies are required to describe the conversion of the ER-born NL lense into the cytoplasmic LD.

In the face of the evidence that the cisterna is fragmented, it is tempting to reject that LDs are mere expansions of the primordial cisterna and that their deflation upon lipolysis reconstitutes the primordial cisterna. For the same reason is more attractive to propose that following lipolysis the LDs materials are recycled into the ER sector specialized in LDs biogenesis. Yet, the fragmentation could be more apparent than real. If real, it can not be excluded the existence of a permanent link between the LDs and the cisterna fragments. Recent studies of the LDs interactome have demonstrated the stability and durability of the LD/ER contacts (Jacquier et al., 2011; Salo et al., 2016; Valm et al., 2017) and the relocation of membrane proteins distributed at the perimeter of LDs into the ER (Blanchette-Mackie et al., 1995; Jacquier et al., 2013).

The possibility that the primordial cisterna is functioning during the period of LD growth is strongly suggested by its appearance in adipocytes with a stable population of large/mature LDs transfected with fluorescent EGFP-E-Syt3ΔC2C, and by the detection of successive waves of new LDs in the maturing adipocyte.

The high-resolution EM and the ET-based 3D reconstruction of the cisterna fragments revealed the existence of spaced units made by a tight network of varicose tubules, wrapped by a membrane, and a cluster of young LDs. Further studies are required to understand the details of this new structure, in particular the connection of the tubules with the adjacent LDs and the origin of the wrapping membrane. The beaded morphology of the tubular membranes suggests a relationship with the known formation of NL droplets between the leaflets of the ER membrane. This interpretation deserves futher study.

Whereas in non-adipose cells the LDs are scattered throughout the cytoplasm in adipocytes, the primordial cisterna appears to fix the spatial localization of the foci from which the LDs bud. In this respect, the retention of the circular profile of the cisterna by the grown LDs is remarkable. It would be interesting to determine if this arrangement facilitates gradual adaptation of the adipocyte to the progressive growth of the droplets and the formation of the single giant droplet that occupies the entire cytoplasm. Given the role of E-Syts as ER/PM tethers and lipid transporters, and the presence of E-Syt3 in the membrane of the single/giant LD in the mature adipocyte, it would be of interest to determine whether E-Syt3 plays a role in fastening the giant LD to the apposed PM and in the exchange of lipids between the two membranes.

E-Syts using the SMP domain to transfer GPLs between the membranes that they bridge (Schauder et al., 2014) and the specific and strong binding of PE and PC to the E-Syt3 SMP domain provide a reasonable background to discuss the results of the effects of E-Syt3 repression on LD biogenesis and growth. The induction of 3T3-L1 fibroblast differentiation into adipocytes increases the conversion of PS into PE in the MT, and PE is then transported to the ER and to the surface of LDs (Horl et al., 2011). These results and the opposite effect of PE and PC on the membrane surface tension suggest that PE is used in the ER to stimulate LD biogenesis, whereas the conversion of PE into PC on the surface of LDs is essential for LD growth and stability (Ben M’barek et al., 2017; Horl et al., 2011). Notably, young adipocytes subjected to repression of E-Syt3 expression exhibited the compact-broad ER collars typically seen before the formation of the first LDs, a phenotype expected in cells with strong inhibition of LD biogenesis. Furthermore, the initial accumulation of small LDs and later growth upon lifting E-Syt3 repression can be explained by the expected increase in PE in the ER and its subsequent conversion into PC. Therefore, it would be important to study whether E-Syt3 is involved in the transport of PE from the MT to the ER and LDs. Thus, it is interesting that ORP5 localizes to MT-ER contact sites and participates in the transfer of PS, the PE precursor, from the ER to MT (Rochin et al., 2019). ORP5 is a membrane protein that also localizes to ER/PM junctions and binds PI(4,5)P2 and, in non-adipose cells, targets LDs and regulates their level of PI(4)P (Du et al., 2020; Ghai et al., 2017; Renne and Emerling, 2020). Advancements in identifying and localizing lipid transporters in the adipocyte and in establishing their possible participation in the formation of permanent and temporal contacts between the MT/ER/LD triad could be decisive in furthering our knowledge of how the exchange of lipids is regulated and its effects on the control of LD biogenesis and growth.

By tracing proteolytic processing and changes in the localization of E-Syt3 and E-Syt3ΔC2C in 3T3-L1 adipocytes, we have identified a new, single, giant cisterna embedded in the ER that is the origin of the formation and growth of LDs. The primordial cisterna, as we call it, is made up of a network of varicose tubules that may provide optimal conditions for the assembly of LDs in adipocytes. The proteasome-regulated C-cleavage of E-Syt3 is essential to targeting the protein to the surface of LDs via a route that includes the primordial cisterna. Repression of E-Syt3 strongly inhibits LD biogenesis and growth. The functioning of E-Syt3 as a GPL transporter on the surface of LDs could be critical for the supply of PE and PC to LDs and, therefore, for LD biogenesis and growth.

## Materials and Methods

### Adipocyte differentiation

Mouse 3T3-L1 fibroblasts maintained at confluence for 48 h in growth medium (DMEM, 1.5 g/L bicarbonate/glutamine/penicillin/streptomycin/10% calf bovine serum) were incubated on days 1 and 2 in adipogenic medium (50 μM 3-isobutyl-1-methylxanthine, 400 nM insulin, 1.5 mM biotin, 0.25 mM dexamethasone, 7.5 mM troglitazone) and on days 3 and 4 with 100 mM insulin. The resulting early adipocytes were kept in plain growth medium that was refreshed every other day to produce mature adipocytes.

### cDNA constructs

*EGFP-tagged E-Syt3 cDNA constructs*. Mouse E-Syt3 cDNA was amplified by PCR from a cDNA library constructed with poly(A) RNA from 3T3-L1 adipocytes (Mate & Plate TM Library System, Clontech) and cloned into the PT-Adv vector using 5’and 3’ E-Syt3 primers (Table 1). Next, E-Syt3 was cloned into the HindIII and Sal I sites of the pEGFP-N2 and EGFP-C2 vectors using specific primers (Table 1). E-Syt3ΔC2C was produced by introducing a 2021_2257 deletion using the Q5 site-directed mutagenesis kit (NEN Biolabs). The deletion created a reading frameshift and a premature stop that removed the C2C domain and created a 707 aa long protein with a 35 aa C-neoextension. When required the C-extension was removed by introducing the substitution 2247 T>A (Table 1). *GST-E-SYT3 constructs*. To generate the N-GST-tagged E-Syt3 fragments SMPC2A, B/C2A, B, C/C2A/C2B and C2C, the E-Syt3 cDNA was amplified using the primers designed for such purpose and the pGEX6P1 vector (Table 1). The 617 aa long E-Syt1ΔC2C, D, E was produced by PCR amplification and cloning into the EGFP-C2 vector using specific primers (Table 1).

**Table 1.**
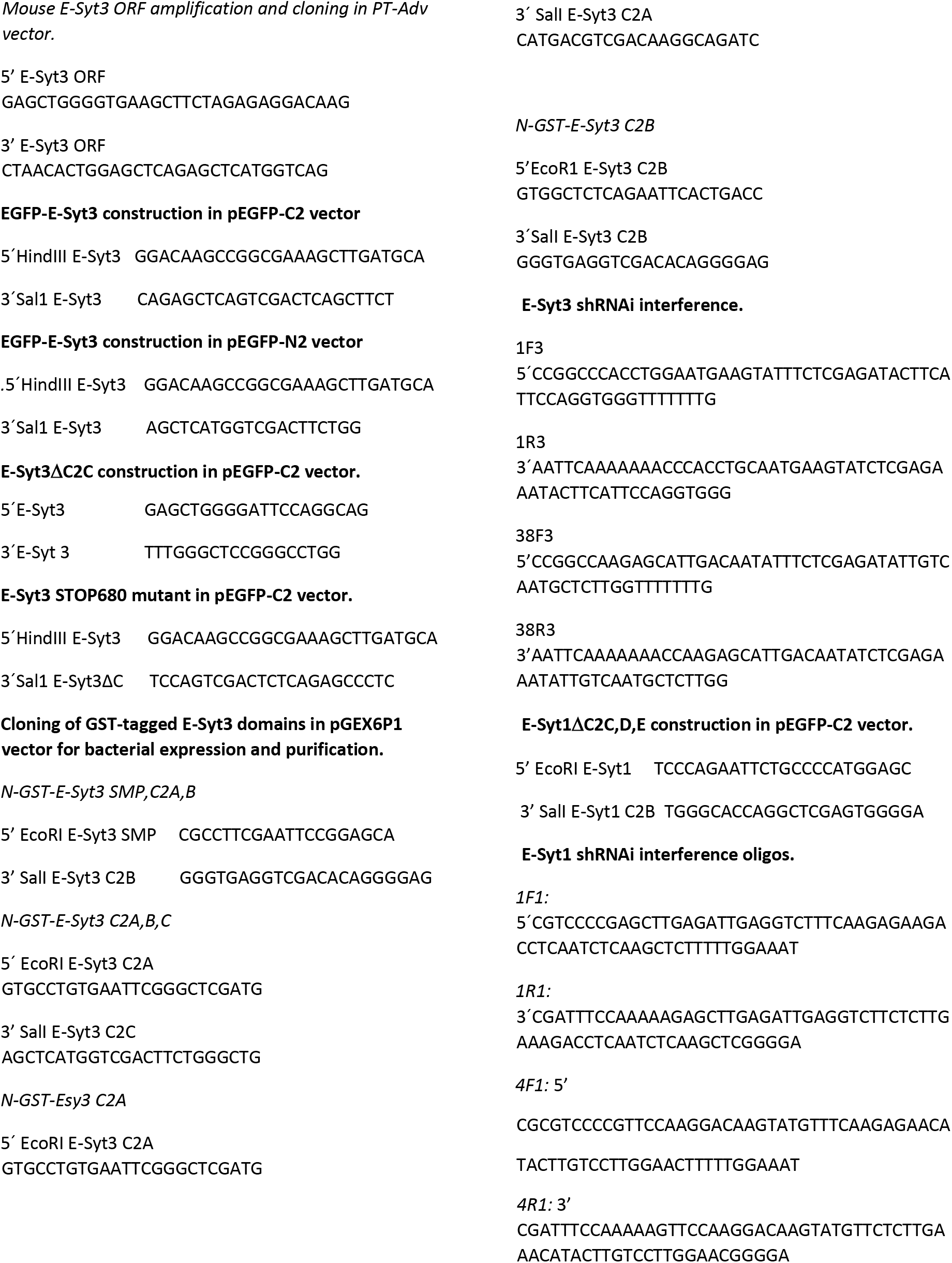
Primers and vectors for DNA modification and amplification.

### RNA interference

The shRNA interference oligos targeting E-Syt3 and E-Syt1 were designed using the N2[CG]N8[AUT]N8 [AUT]N2 rule (Pei and Tuschl, 2006) at siRNA WhiteHead (Table 1) and cloned into the pLKO-Tet-On and pLKO pLKO.1-cherry vectors, respectively (Stewart et al., 2003); (Szulc and Aebischer, 2008). Vector particles were produced from second day supernatants conditioned by 293FT cells transduced using 8 μg/ml polybrene (Invitrogen) (Szulc and Aebischer, 2008). For pLKO-Tet-On shRNA interference, 3T3-L1 fibroblasts at 30% confluence were incubated with lentiviral particles (5 MOIs) and selected with 2.9 μg/ml puromycin, carefully avoiding cell confluency. To induce shRNA expression and gene knockdown, the puromycin selected 3T3-L1 fibroblasts were grown in 200 ng/ml doxycycline (Dox) for different time periods as required. For pLKO.1-cherry shRNA interference, 3T3-L1 fibroblasts were infected with lentiviral particles, selected with 2.9 μg/ml puromycin and separated using a fluorescence-activated cell sorter (FACS).

### 3T3-L1 adipocyte transfection

3T3-L1 adipocytes (2×10^6^ cells) on day 4 of differentiation were harvested by 10 min of trypsinization, collected by low speed centrifugation, carefully resuspended in 100 μl Cell Line Nucleofector Solution L containing 2 μg DNA and electroporated using an Amaxa Nucleofector set to program A-033. Confluent 3T3-L1 fibroblasts (2×10^6^ cells) were electroporated using Cell Line Nucleofector Solution V and program T-030. After their plating on plastic dishes or cover glasses the cells were used for microcopy and biochemical studies as indicated in Results.

### Confocal immunofluorescence microscopy

Cells grown on 10 mm glass coverslips were fixed-permeabilized using conditions adapted to the reactivity of the antibodies. The cells were quickly washed in PBS at room temperature and fixed-permeabilized for 4 min with cold (−20°C) methanol. Alternatively, fixation was performed with 2% paraformaldehyde/PBS for 10 min, permeabilization was performed with 0.2% Triton X-100/PBS for 5 min and the reactive aldehydes were blocked with 10 mM glycine for 20 min; all the protocol steps were performed at room temperature. Proteins were stained using specific antibodies, neutral lipids were stained with HCS LipidTOX red neutral (Thermo Fisher) and DNA was stained using DAPI. Glass coverslips were mounted on Gelvatol (refractive index 1.376). Stable Alexa 488, Alexa 555, Alexa 647 and Pacific Blue dyes were used to avoid spectral bleeding. The stained cells were examined using Zeiss Plan-Neofluar x40 and Plan-apochromat x63/1.4a oil DIC objectives and oil (r.i. 1.51). Immunofluorescence microscopy (IFM) images were uploaded from a confocal Zeiss Spectral Multifoton LSM710. Protein colocalization was studied using a confocal multispectral Leica TCS SP8 system equipped with a 3X STED module for super-resolution (laser line: 405 nm. depletion line 660 nm). Confocal images were studied using the ImajeJ 1.52o/t platforms. Image deconvolution was performed using the Huygens deconvolution software.

### Live-cell time-lapse fluorescence and bright-field microscopy

Cell imaging acquisition was performed using a Zeiss Axiovert 200 microscope equipped with a Zeiss x40 oil immersion objective and a Hamamatsu ORCA-Flash 4.9 LT sCMOS digital camera. Metamorph 7.10 software was used for automatic image acquisition, equipment coordination and image analysis. The cellular distribution of E-Syt3 was studied on images captured every 10 min in seven separate planes over 13-22 h. LD fusion was studied by analyzing the object number, size and average area on watershed segmented images after setting an Auto Local Threshold with the Bemsem method/radius 5.

### Correlative-light immunoelectron microscopic analysis (CLEM)

Fluorescence microscopy was combined with immune EM and three-dimensional correlative electron tomography (Micaroni et al., 2010) (Fusella et al., 2013) (Beznoussenko et al., 2016) to study the ultrastructure of specific segments of type 1 and type 2 primordial cisternae and their association with E-Syt3ΔC2C. The necessary cell fixation with 4% paraformaldehyde and 0.05% glutaraldehyde precluded the use of the anti-E-Syt3 antibodies. Accordingly, pre-embedding immune EM detection of E-Syt3ΔC2C was performed in young adipocytes on day 3 of differentiation using rabbit anti-EGFP antibody and goat anti-rabbit Fab’ fragments coupled to 1.4 nm gold particles as described (Kweon et al., 2004) (Beznoussenko et al., 2015). TEM tomograms were reconstructed by acquiring images automatically at a series of specimen tilt using the iterative reconstruction technique implemented by the program Xplore3DTMXpress-Inspect3DTM and the last version of the IMOD package (Mastronarde, 2005).

### Antibodies

Polyclonal α-E-Syt3p^1^ antibody was developed in rabbit and rat against the E-Syt3 peptides C-141ENKIREKLEPKIREKS^156^, C-^371^EVPGQDLEVDLYDEDTDKDAD^390^ and C-^850^SRPLGSHRRKELGK^863^ (see Fig. S2) and purified by affinity chromatography using the immunized serum and the three antigenic peptides bound to Sulfolink resin to study the cellular distribution of E-Syt3. The α-E-Syt3p^141^ antibody was purified by affinity chromatography using α-E-Syt3p^1^ serum and peptide^141-156^ bound to Sulfolink resin and used to study the proteolytic processing of the N- and C-EGFP tagged E-Syt3 constructs. Antibody α-E-Syt3p^77^ was raised in rat using the peptide^77^RRNRRGKLGRLEAAFE FLEHEREFISRELRGQH^109^ and used to study the distribution of E-Syt3 in 3T3-L1 fibroblasts and young adipocytes. Polyclonal α-E-Syt1 antibody was developed in rabbit against the peptides C-1MEHSPEEGASPEPSGQ^16^, C-^1038^TLRRKLDVSVKSN^1052^, C-^1038^TLRRKLDVSVKSNSS^1020^ and was affinity purified using the antigenic peptides.

Other antibodies were, mouse monoclonal antibodies targeting GFP/EGFP (Roche), ubiquitin P4G7 (Biolegend), lysosomal Lamp I (Barriocanal et al., 1986), Na^+^/K^+^-ATPase alpha-1 subunit 2F (DSHB Iowa University), lamin B2 LN43 8983 (Abcam), β-catenin 610154 (BD Transduction), actin 224-236-1 (DSHB Iowa University) and GST clone 6G9C6 (Sigma-Aldrich); rabbit polyclonal anti-GFP/EGFP PABG1 (Chromotek), anti-calnexin (StressMarq Biosciences) and PDI (Dr. José Gonzalez Castaño, UAM) antibodies; goat polyclonal anti-PLIN1 ab61682 (Abcam) antibody; and guinea pig anti-PLIN-2 and PLIN3 antibodies (Progen Biotech. GP46, GP30). The secondary antibodies used were Alexa 488, Alexa 555 and Alexa 647, Pacific Blue (Life Technologies), horseradish peroxidase (HRP)-conjugated IgG F(ab)2 donkey anti-rabbit (GE Healthcare), and HRP-conjugated affinity-purified goat anti-mouse IgG (Jackson).

### Preparation of adipocyte extracts, subcellular fractionation, LD purification and immunoblotting

All the manipulations were performed at 4°C. 3T3-L1 adipocytes were washed twice with PBS, scrapped in cold hypotonic buffer A (10 mM HEPES at pH 7.5 with 0.25 M sucrose, 1 mM EDTA, 50 mM sodium fluoride, 1 mM β-glycerol phosphate, 1 mM orthovanadate and the cocktail of protease inhibitors cOmplete Mini, Roche) and collected by centrifugation for 5 min at 500xg. For subcellular fractionation, the cells were disrupted by 20 up and down passages through a 23 gauge needle, and the extracts were centrifuged at 1000xg for 10 min to obtain a crude nuclear and a postnuclear fraction (PN). Purified nuclei (pN) were prepared from crude nuclear pellets resuspended in buffer B (10 mM HEPES at pH 7.9 with 0.25 M sucrose, 10 mM KCl, 1.5 mM MgCl_2_, mM DTT) containing the phosphatases and proteases inhibitors and centrifuged at 100,000xg for 1 h through a 3 ml 1.4 M sucrose cushion using a TL100 centrifuge (Beckman). The crude heavy nuclear fraction and purified nuclei were resuspended in buffer A and extracted with 1% Triton X-100 for 5 min at 4°C (cNM, pNM). LDs floating on postnuclear supernatants were purified by sucrose step-gradient centrifugation, essentially as described (Brasaemle et al., 2004). For protein analysis purified LDs suspended in hypotonic buffer were mixed with 1 volume of 1.5% Triton X-114 and then laid over a 6% sucrose cushion and incubated for 3 min at 30°C to concentrate the membrane proteins by centrifugation for 5 min at 5000xg. Protein delipidation was performed as described (Wessel and Flugge, 1984). Plasma membrane sheets were purified from the PN and NM fractions, respectively (Simpson et al., 1983). For analysis in the cellular fractions, the proteins were resolved by SDS-polyacrylamide gel electrophoresis and studied by immunoblotting and chemiluminescence using HRP-conjugated secondary antibodies. To study the proteolytic products of the transfected EGFP-tagged E-Syt constructs, the amount of protein and the concentration of the primary antibodies were carefully titrated to minimize interference with the endogenous E-Syt3 species (Fig. 2).

### Binding of PLs to E-Syt3 domains

BL21(DE3)pLysS competent cells (Promega) transformed with N-GSTE-Syt3 domain constructs carried in the pGEX6P1 vector were resuspended in STE buffer (150 mM NaCl, 10 mM Tris-HCl at pH 8.0, 1 mM EDTA) and incubated with 50 μg/ml lysozyme for 15 min on ice. Then, DTT was added to 1 mM, and N-laurylsarcosine was added to 1.5%. After 3 sonication steps for 40 sec each, the cell extracts were centrifuged to 10,000xg for 10 min, and the supernatants were treated with 1% NP-40 for 30 min at room temperature. Following the removal of insoluble material, the detergent extracts were incubated with glutathione Sepharose 4B (Amersham Biosciences) for 2 h at 4°C to batch purify the GST recombinant constructs. The beads were extensively washed with STE buffer and eluted with 0.1 M glycine at pH 2.5. Next, the pH was immediately neutralized with 1 M Tris base. The purified recombinant C2ABC, SMPC2AB, C2A, C2B and C2C domains (0.5 μg/ml) were incubated for overnight at 4°C with PIP strips (Echelon Biosciences) containing 100 pmol of 15 PL points distributed in nitrocellulose strips. The binding of the GST constructs to PLs was studied by immunoblotting using a polyclonal anti-GST antibody in conjunction with a horseradish peroxidase secondary antibody and was quantified by chemiluminescence.

## Acknowledgments

The authors expressed their gratitude to Dr. David Abia for help with the E-Syt3 modeling, to Ms. Sylvia Gutierrez, Dr. Ana Oña and Ms. Angeles Muñoz for help with the high-resolution confocal microscopy and Dr. Carmen Sanchez and Mr. José Ignacio Belio for help with the videos and art-work. This work was supported by grants from the Instituto de Salud Carlos III (grant number PI17/02277) and the Fundación Areces.

## Author Contributions

V.L. and I.V.S. planned the use and produced the biological tools (cDNA constructs and anti-E-Syt antibodies), performed the cell biology and biochemical experiments and wrote the manuscript. G.B and A. M performed the EM and ET studies.

## References

Barriocanal, J. G., Bonifacino, J.S., Yuan, L., and Sandoval, I.V. (1986). Biosynthesis, glycosylation, movement through the Golgi system, and transport to lysosomes by an N-linked carbohydrate-independent mechanism of three lysosomal integral membrane proteins. J Biol Chem 261, 16755–16763.

Ben M’barek, K., Ajjaji, D., Chorlay, A., Vanni, S., Foret, L., and Thiam, A.R. (2017). ER Membrane Phospholipids and Surface Tension Control Cellular Lipid Droplet Formation. Dev Cell 41, 591–604 e597.

Beznoussenko, G.V., Pilyugin, S.S., Geerts, W.J., Kozlov, M.M., Burger, K.N., Luini, A., Derganc, J., and Mironov, A.A. (2015). Trans-membrane area asymmetry controls the shape of cellular organelles. Int J Mol Sci 16, 5299–5333.

Beznoussenko, G.V., Ragnini-Wilson, A., Wilson, C., and Mironov, A.A. (2016). Three-dimensional and immune electron microscopic analysis of the secretory pathway in Saccharomyces cerevisiae. Histochem Cell Biol 146, 515–527.

Blanchette-Mackie, E.J., Dwyer, N.K., Barber, T., Coxey, R.A., Takeda, T., Rondinone, C.M., Theodorakis, J.L., Greenberg, A.S., and Londos, C. (1995). Perilipin is located on the surface layer of intracellular lipid droplets in adipocytes. J Lipid Res 36, 1211–1226.

Brasaemle, D.L., Barber, T., Kimmel, A.R., and Londos, C. (1997). Post-translational regulation of perilipin expression. Stabilization by stored intracellular neutral lipids. J Biol Chem 272, 9378–9387.

Brasaemle, D.L., Dolios, G., Shapiro, L., and Wang, R. (2004). Proteomic analysis of proteins associated with lipid droplets of basal and lipolytically stimulated 3T3-L1 adipocytes. J Biol Chem 279, 46835–46842.

Choudhary, V., Ojha, N., Golden, A., and Prinz, W.A. (2015). A conserved family of proteins facilitates nascent lipid droplet budding from the ER. J Cell Biol 211, 261–271.

Chung, J., Wu, X., Lambert, T.J., Lai, Z.W., Walther, T.C., and Farese, R.V.Jr. (2019). LDAF1 and Seipin Form a Lipid Droplet Assembly Complex. Dev Cell 51, 551–563 e557.

Du, X., Zhou, L., Aw, Y.C., Mak, H.Y., Xu, Y., Rae, J., Wang, W., Zadoorian, A., Hancock, S.E., Osborne, B., et al. (2020). ORP5 localizes to ER-lipid droplet contacts and regulates the level of PI(4)P on lipid droplets. J Cell Biol 219.

Fusella, A., Micaroni, M., Di Giandomenico, D., Mironov, A.A., and Beznoussenko, G.V. (2013). Segregation of the Qb-SNAREs GS27 and GS28 into Golgi vesicles regulates intra-Golgi transport. Traffic 14, 568–584.

Ghai, R., Du, X., Wang, H., Dong, J., Ferguson, C., Brown, A.J., Parton, R.G., Wu, J.W., and Yang, H. (2017). ORP5 and ORP8 bind phosphatidylinositol-4, 5-biphosphate (PtdIns(4,5)P 2) and regulate its level at the plasma membrane. Nat Commun 8, 757.

Giordano, F., Saheki, Y., Idevall-Hagren, O., Colombo, S.F., Pirruccello, M., Milosevic, I., Gracheva, E.O., Bagriantsev, S.N., Borgese, N., and De Camilli, P. (2013). PI(4,5)P(2)-dependent and Ca(2+)-regulated ER-PM interactions mediated by the extended synaptotagmins. Cell 153, 1494–1509.

Haemmerle, G., Moustafa, T., Woelkart, G., Buttner, S., Schmidt, A., van de Weijer, T., Hesselink, M., Jaeger, D., Kienesberger, P.C., Zierler, K., et al. (2011). ATGL-mediated fat catabolism regulates cardiac mitochondrial function via PPAR-alpha and PGC-1. Nat Med 17, 1076–1085.

Heid, H.W., Moll, R., Schwetlick, I., Rackwitz, H.R., and Keenan, T.W. (1998). Adipophilin is a specific marker of lipid accumulation in diverse cell types and diseases. Cell Tissue Res 294, 309–321.

Henne, M., Goodman, J.M., and Hariri, H. (2019). Spatial compartmentalization of lipid droplet biogenesis. Biochim Biophys Acta Mol Cell Biol Lipids.

Herdman, C., and Moss, T. (2016). Extended-Synaptotagmins (E-Syts); the extended story. Pharmacol Res 107, 48–56.

Herdman, C., Tremblay, M.G., Mishra, P.K., and Moss, T. (2014). Loss of Extended Synaptotagmins ESyt2 and ESyt3 does not affect mouse development or viability, but in vitro cell migration and survival under stress are affected. Cell Cycle 13, 2616–2625.

Horl, G., Wagner, A., Cole, L.K., Malli, R., Reicher, H., Kotzbeck, P., Kofeler, H., Hofler, G., Frank, S., Bogner-Strauss, J.G., et al. (2011). Sequential synthesis and methylation of phosphatidylethanolamine promote lipid droplet biosynthesis and stability in tissue culture and in vivo. J Biol Chem 286, 17338–17350.

Itabe, H., Yamaguchi, T., Nimura, S., and Sasabe, N. (2017). Perilipins: a diversity of intracellular lipid droplet proteins. Lipids Health Dis 16, 83.

Jacquier, N., Choudhary, V., Mari, M., Toulmay, A., Reggiori, F., and Schneiter, R. (2011). Lipid droplets are functionally connected to the endoplasmic reticulum in Saccharomyces cerevisiae. J Cell Sci 124, 2424–2437.

Jacquier, N., Mishra, S., Choudhary, V., and Schneiter, R. (2013). Expression of oleosin and perilipins in yeast promotes formation of lipid droplets from the endoplasmic reticulum. J Cell Sci 126, 5198–5209.

Jiang, H.P., and Serrero, G. (1992). Isolation and characterization of a full-length cDNA coding for an adipose differentiation-related protein. Proc Natl Acad Sci U S A 89, 7856–7860.

Kassan, A., Herms, A., Fernandez-Vidal, A., Bosch, M., Schieber, N.L., Reddy, B.J., Fajardo, A., Gelabert-Baldrich, M., Tebar, F., Enrich, C., et al. (2013). Acyl-CoA synthetase 3 promotes lipid droplet biogenesis in ER microdomains. J Cell Biol 203, 985–1001.

King, N.J., Delikatny, E.J., and Holmes, K.T. (1994). 1H magnetic resonance spectroscopy of primary human and murine cells of the myeloid lineage. Immunomethods 4, 188–198.

Krahmer, N., Guo, Y., Wilfling, F., Hilger, M., Lingrell, S., Heger, K., Newman, H.W., Schmidt-Supprian, M., Vance, D.E., Mann, M., et al. (2011). Phosphatidylcholine synthesis for lipid droplet expansion is mediated by localized activation of CTP:phosphocholine cytidylyltransferase. Cell Metab 14, 504–515.

Kweon, H.S., Beznoussenko, G.V., Micaroni, M., Polishchuk, R.S., Trucco, A., Martella, O., Di Giandomenico, D., Marra, P., Fusella, A., Di Pentima, A., et al. (2004). Golgi enzymes are enriched in perforated zones of golgi cisternae but are depleted in COPI vesicles. Mol Biol Cell 15, 4710–4724.

Lalioti, V., Muruais, G., Dinarina, A., van Damme, J., Vandekerckhove, J., and Sandoval, I.V. (2009). The atypical kinase Cdk5 is activated by insulin, regulates the association between GLUT4 and E-Syt1, and modulates glucose transport in 3T3-L1 adipocytes. Proc Natl Acad Sci U S A 106, 4249–4253.

Li, Z., Thiel, K., Thul, P.J., Beller, M., Kuhnlein, R.P., and Welte, M.A. (2012). Lipid droplets control the maternal histone supply of Drosophila embryos. Curr Biol 22, 2104–2113.

Mackinnon, W.B., May, G.L., and Mountford, C.E. (1992). Esterified cholesterol and triglyceride are present in plasma membranes of Chinese hamster ovary cells. Eur J Biochem 205, 827–839.

Manford, A.G., Stefan, C.J., Yuan, H.L., Macgurn, J.A., and Emr, S.D. (2012). ER-to-plasma membrane tethering proteins regulate cell signaling and ER morphology. Dev Cell 23, 1129–1140.

Martin, S., and Parton, R.G. (2006). Lipid droplets: a unified view of a dynamic organelle. Nat Rev Mol Cell Biol 7, 373–378.

Mastronarde, D.N. (2005). Automated electron microscope tomography using robust prediction of specimen movements. J Struct Biol 152, 36–51.

Micaroni, M., Perinetti, G., Berrie, C.P., and Mironov, A.A. (2010). The SPCA1 Ca2+ pump and intracellular membrane trafficking. Traffic 11, 1315–1333.

Min, S.W., Chang, W.P., and Sudhof, T.C. (2007). E-Syts, a family of membranous Ca2+-sensor proteins with multiple C2 domains. Proc Natl Acad Sci U S A 104, 3823–3828.

Mishra, S., Khaddaj, R., Cottier, S., Stradalova, V., Jacob, C., and Schneiter, R. (2016). Mature lipid droplets are accessible to ER luminal proteins. J Cell Sci 129, 3803–3815.

Murphy, D.J., and Vance, J. (1999). Mechanisms of lipid-body formation. Trends Biochem Sci 24, 109–115.

Ohsaki, Y., Soltysik, K., and Fujimoto, T. (2017). The Lipid Droplet and the Endoplasmic Reticulum. Adv Exp Med Biol 997, 111–120.

Pei, Y., and Tuschl, T. (2006). On the art of identifying effective and specific siRNAs. Nat Methods 3, 670–676.

Renne, M.F., and Emerling, B.M. (2020). ORP5 regulates PI(4)P on the lipid droplet: Novel players on the monolayer. J Cell Biol 219.

Rochin, L., Sauvanet, C., Jääskeläinen, E., Houcine, A., Kivelä, A., MA, X., Marien, E., Dehairs, J., Neveu, J., Bars, R.L., et al. (2019). ORP5 transfers phosphatidylserine to mitochondria and regulates mitochondrial calcium uptake at endoplasmic reticulum-mitochondria contact sites. bioRxiv Posted.

Saheki, Y., Bian, X., Schauder, C.M., Sawaki, Y., Surma, M.A., Klose, C., Pincet, F., Reinisch, K.M., and De Camilli, P. (2016). Control of plasma membrane lipid homeostasis by the extended synaptotagmins. Nat Cell Biol 18, 504–515.

Saka, H.A., and Valdivia, R. (2012). Emerging roles for lipid droplets in immunity and host-pathogen interactions. Annu Rev Cell Dev Biol 28, 411–437.

Salo, V.T., Belevich, I., Li, S., Karhinen, L., Vihinen, H., Vigouroux, C., Magre, J., Thiele, C., Holtta-Vuori, M., Jokitalo, E., et al. (2016). Seipin regulates ER-lipid droplet contacts and cargo delivery. EMBO J 35, 2699–2716.

Schauder, C.M., Wu, X., Saheki, Y., Narayanaswamy, P., Torta, F., Wenk, M.R., De Camilli, P., and Reinisch, K.M. (2014). Structure of a lipid-bound extended synaptotagmin indicates a role in lipid transfer. Nature 510, 552–555.

Simpson, I.A., Yver, D.R., Hissin, P.J., Wardzala, L.J., Karnieli, E., Salans, L.B., and Cushman, S.W. (1983). Insulin-stimulated translocation of glucose transporters in the isolated rat adipose cells: characterization of subcellular fractions. Biochim Biophys Acta 763, 393–407.

Stein, O., and Stein, Y. (1967). Lipid synthesis, intracellular transport, and secretion. II. Electron microscopic radioautographic study of the mouse lactating mammary gland. J Cell Biol 34, 251–263.

Stewart, S.A., Dykxhoorn, D.M., Palliser, D., Mizuno, H., Yu, E.Y., An, D.S., Sabatini, D.M., Chen, I.S., Hahn, W.C., Sharp, P.A., et al. (2003). Lentivirus-delivered stable gene silencing by RNAi in primary cells. RNA 9, 493–501.

Szulc, J., and Aebischer, P. (2008). Conditional gene expression and knockdown using lentivirus vectors encoding shRNA. Methods Mol Biol 434, 291–309.

Verma, R., and Deshaies, R.J. (2000). A proteasome howdunit: the case of the missing signal. Cell 101, 341–344.

Walther, T.C., Chung, J., and Farese, R.V. Jr. (2017). Lipid Droplet Biogenesis. Annu Rev Cell Dev Biol 33, 491–510.

Walther, T.C., and Farese, R.V. Jr. (2012). Lipid droplets and cellular lipid metabolism. Annu Rev Biochem 81, 687–714.

Wang, H., Becuwe, M., Housden, B.E., Chitraju, C., Porras, A.J., Graham, M.M., Liu, X.N., Thiam, A.R., Savage, D.B., Agarwal, A.K., et al. (2016). Seipin is required for converting nascent to mature lipid droplets. Elife 5.

Wang, S., Idrissi, F.Z., Hermansson, M., Grippa, A., Ejsing, C.S., and Carvalho, P. (2018). Seipin and the membrane-shaping protein Pex30 cooperate in organelle budding from the endoplasmic reticulum. Nat Commun 9, 2939.

Wanner, G., Formanek, H., and Theimer, R.R. (1981). The ontogeny of lipid bodies (spherosomes) in plant cells: Ultrastructural evidence. Planta 151, 109–123.

Wessel, D., and Flugge, U.I. (1984). A method for the quantitative recovery of protein in dilute solution in the presence of detergents and lipids. Anal Biochem 138, 141–143.

