## Supplemental Materials for "The Step-Wise C-Truncation and Transport of ESyt3 to Lipid Droplets Reveals a Mother Primordial Cisterna"

**Videos.**

Videos can be reached at https://babia.cbm.uam.es:5001/sharing/Pbl5eXNT2 Open and play the videos using ImageJ.

**Videos 1 and 2.**

**Live-cell time-lapse fluorescence microscopy of EGFP-E-Syt3ΔC2C transit through the primordial cisterna from which LDs bud.** Adipocytes were transfected with EGFP-E-Syt3ΔC2C on day 4 of differentiation. Beginning 3 h after transfection, the dynamics of the subcellular redistribution of the protein were recorded in 10 separate fields every 10 min and the best focus among the seven recorded planes was used to mount the videos. Note the segmentation of the fluorescent cisterna immediately before the formation of the first LDs: video 1 (00.40-01.30 h), video 2 (11.30-11.50h).

**Videos 3 and 4.**

**EGFP-E-Syt3ΔC2C expression and relocation during the early stages of LD assembly and growth.** Adipocytes were studied on day 4 of differentiation as described in videos 1 and 2. Neutral lipids (NLs) were stained with red LipidTOX. Bright field and fluorescent images were recorded sequentially. Note the four consecutive phases of EGFP-E-Syt3ΔC2C distribution from the initial diffuse cytoplasmic distribution (00:00) and its progressive accumulation into the cortical ER (02:40) to its concentration into discrete cisterna segments (04:20) and the appearance of NL microdrops in their lumen (10:10-16:50). Also, note the fusion between the cisternae containing microdrops (see Fig. 6A, B).

**Videos 5 and 6.**

**Live-cell study of LD homotypic fusion.**

Control adipocytes stained with LipidTOX on day 4 of differentiation (video 5) and adipocytes transfected with EGFP-E-Syt3ΔC2C on day 4 (video 6). Images were recorded every 10 min. Note the surface decoration of LDs with EGFP-E-Syt3ΔC2C (i.e., video 7, t 19:10-20:40) and the predation of small LDs by large LDs.

**Video 7.**

**Control experiment showing the stable retention of the ER membrane marker Sec61β throughout the ER.**

Adipocytes were transfected with ER marker protein EGFP-Sec61β on day 4 of differentiation. Images were recorded every 10 min. Note the homogenous spread and retention of the fluorescent protein through the entire ER.

All video experiments were repeated at least two times.

### Appendix Fig. S1

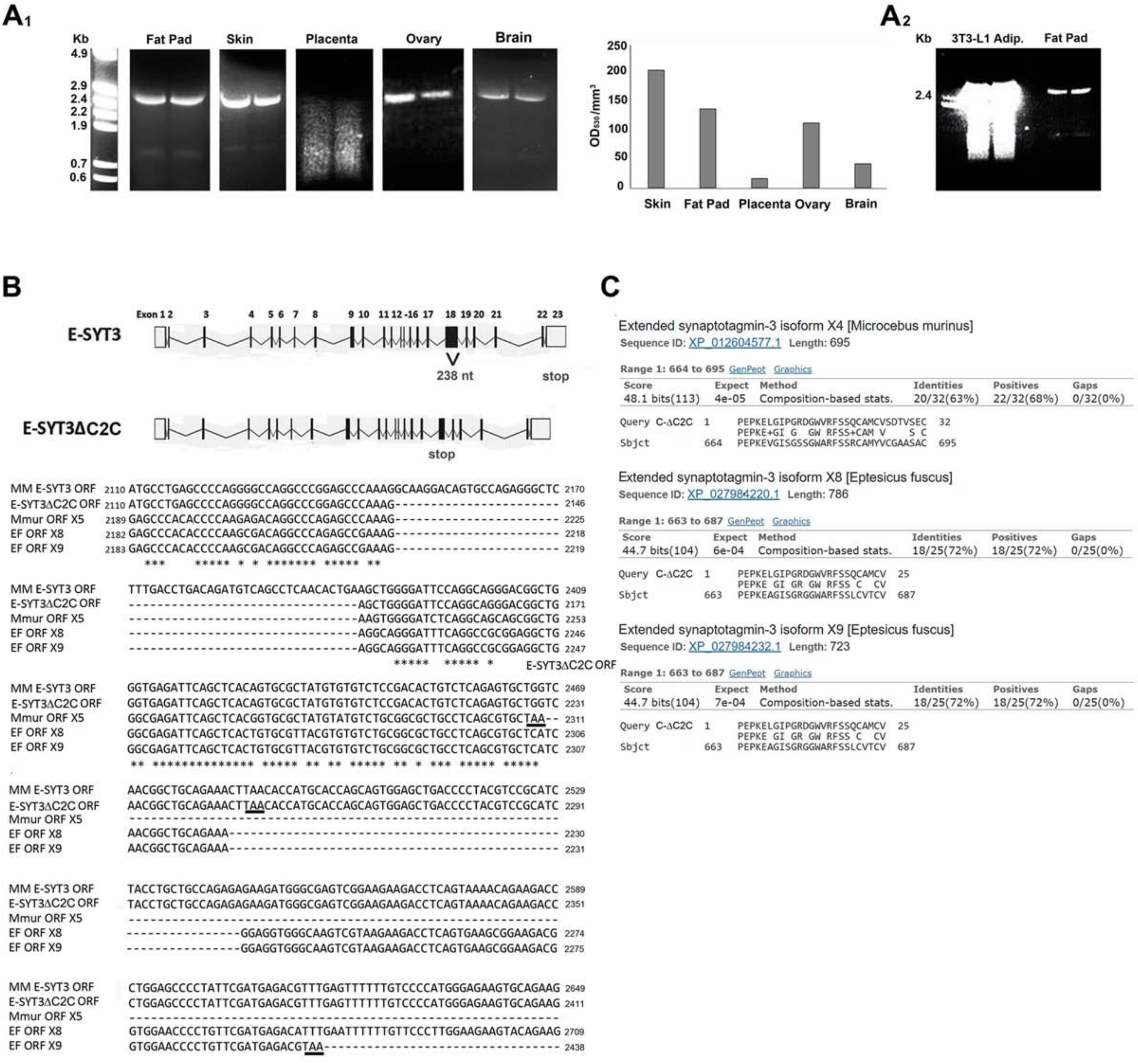


**Fig. S1 E-Syt3 transcripts in mouse tissues and 3T3-L1 adipocytes. E-Syt3□C2C construction.**

($A_1$) Amplification of E-Syt3 transcripts by RT-PCR using 0.1 and 0.05 µg of purified mRNA pretreated with DNAse and 3', 5'-specific E-Syt3 primers (Table 1) resulted in significant levels of the 2.7 kb coding sequence of the E-Syt3 transcript variant X1 (XM_006511221.3) in mouse skin, fat pad, ovary, and brain tissues. ($A_2$) Amplification of E-Syt3 transcripts from 3T3-L1 adipocytes and purified mRNA from mouse fat pad. (B, C) E-Syt3□C2C construction. To delete the C2C domain from mouse E-Syt3, we created a 238 nt deletion in exon 18 (nt 2014- 2252) that mimicked the deletions produced by reading frameshifts and premature stops predicted by shotgun in *Microcebus murinus* and in *Eptesicus fuscus*. Purified mouse 3T3-L1 adipocyte mRNA and specific E-Syt3 mRNA oligos (see Table 1) were used. The resulting E-Syt3□C2C was 707 aa in length, lacked the C2C domain, and ended in a 35 aa C-terminal neosequence homologous to the sequence found in *Microcebus murinus* and *Eptesicus fuscus* (C). The C-neosequence was removed in the construct E-Syt3STOP$^{680}$ (see Table 1, Fig. S6).

### Appendix Fig. S2

A1

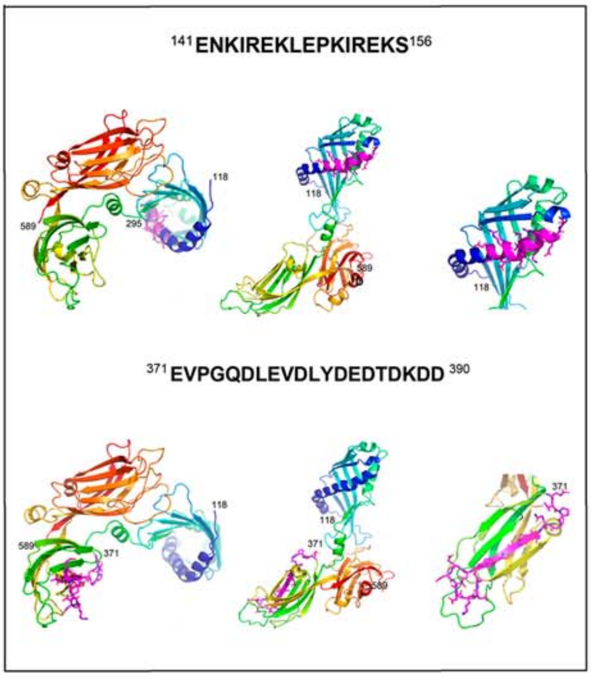


A2

| α-E-Syt3p1 | | OD450 |
|---|---|---|
| peptide 141-156 | 1μg | 0.934 |
| peptide 371-390 | 1μg | 0.464 |
| peptide 850-863 | 1μg | 1.085 |
| E-Syt3 SMP, C2A,B | 1μg | 0.715 |
| E-Syt3 C2A,B,C | 1μg | 0.640 |
| unrelated peptide | 1μg | 0.182 |

A3

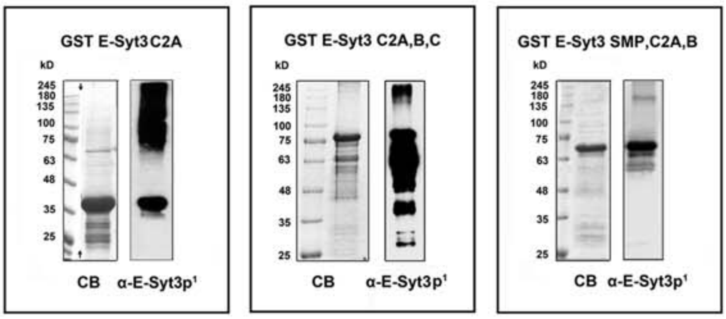


A4

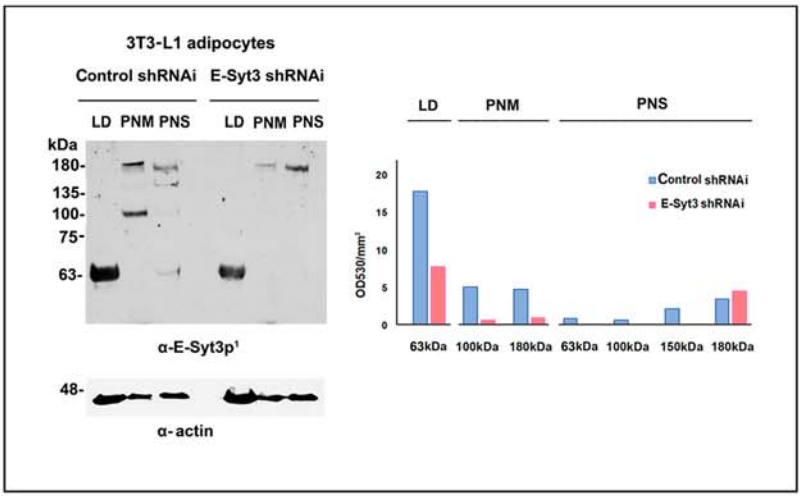


A5

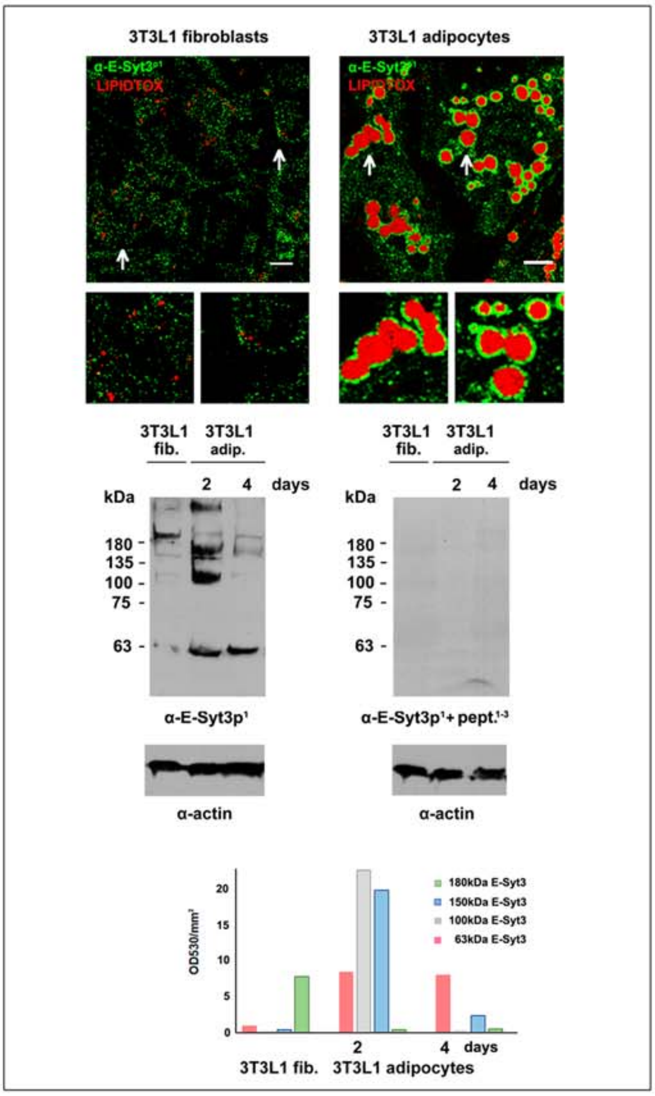


A6

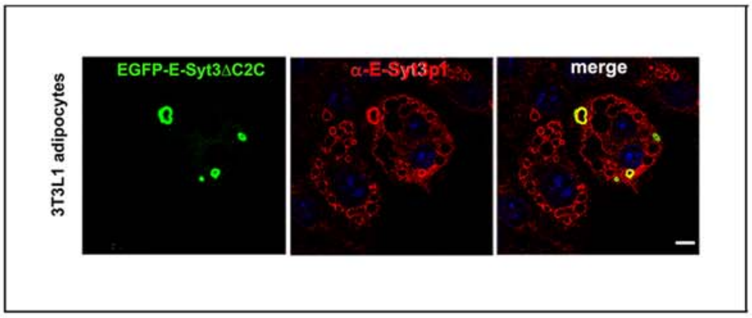

**Fig. S2 Validation of affinity purified rabbit α-E-Syt3 $p^1$**

Polyclonal α-E-Syt3 antibodies were raised in rabbit and rat against the peptides $^{141}$ENKIREKLEPKIREKS$^{156}$, C-$^{371}$EVPGQDLEVDLYDEDTDKDAD$^{390}$, and C-$^{850}$SRPLGSHRRKELGK$^{863}$. The rabbit and rat antibodies produced comparable results in microscopy and Western blot studies. The rabbit α-E-Syt3$p^1$ antibody was used as a source to produce the affinity purified α-E-Syt3$p^{141}$ antibody. ($A_1$) Homology model of the E-Syt3 fragment$^{118\text{-}589}$ constructed according to the template provided by the E-Syt2 SMP,C2A,B crystal (PDB, ID 4p42) (Schauder et al., 2014). Note the exposure of antigenic peptides 141-156 and 371-390 (magenta) on the surface of the SMP (aa 118-295) and C2A (aa 311-411) domains, respectively. Left to right, en face and lateral views. No structural information was available on the domain containing the E-Syt3 peptide$^{850\text{-}863}$. ($A_2$) ELISA assay showing the reactivity of α-E-Syt3$p^1$ with the three antigenic peptides and the purified soluble E-Syt3 SMP,C2A,B and E-Syt3 C2A,B,C fragments; side-reaction with the unrelated LAMP1 KVGNKSRVLELQFGM peptide. ($A_3$) Coomassie blue (CB) and reactivity of antibody α-E-Syt3$p^1$ with soluble GST-conjugated E-Syt3 C2A, C2A, B, C and SMP, C2A, B fragments expressed in bacteria. ($A_4$) Western blot analysis of the α-E-Syt3$p^1$ reactivity with purified LDs (LD, 15 µg), whole postnuclear membranes (PNM, 20 µg), and postnuclear supernatant (PNS, 20 µg) from control 3T3-L1 adipocytes and 3T3-L1 adipocytes subjected to E-Syt3 shRNA repression for 12 days. Actin was used as a loading control. Note the specific reaction of the antibody with the 180 kDa, 150 kDa, 100 kDa, and 63 kDa E-Syt3 species and the species repression by E-Syt3 shRNA interference. ($A_5$) Confocal IFM. Observe the decoration of the LDs surface with antibody α-E-Syt3$p^1$ in the 3T3-L1 adipocyte as well as the inhibiton of the antibody reactivity with the major E-Syt3 species in extracts from 3T3-L1 fibroblasts and 3T3-L1 adipocytes after preincuabtion with the antigenic peptides (pept.$^{1\text{-}3}$). ($A_6$) 3T3-L1 adipocytes were transfected with EGFP-E-Syt3ΔC2C. Note the proportional reaction of the antibody with the endogenous and overexpressed protein. Scale bars = 10 µm. The ELISA and western studies described were repeated twice and the IFM at least three times with comparable results.

#### Appendix Fig.S3

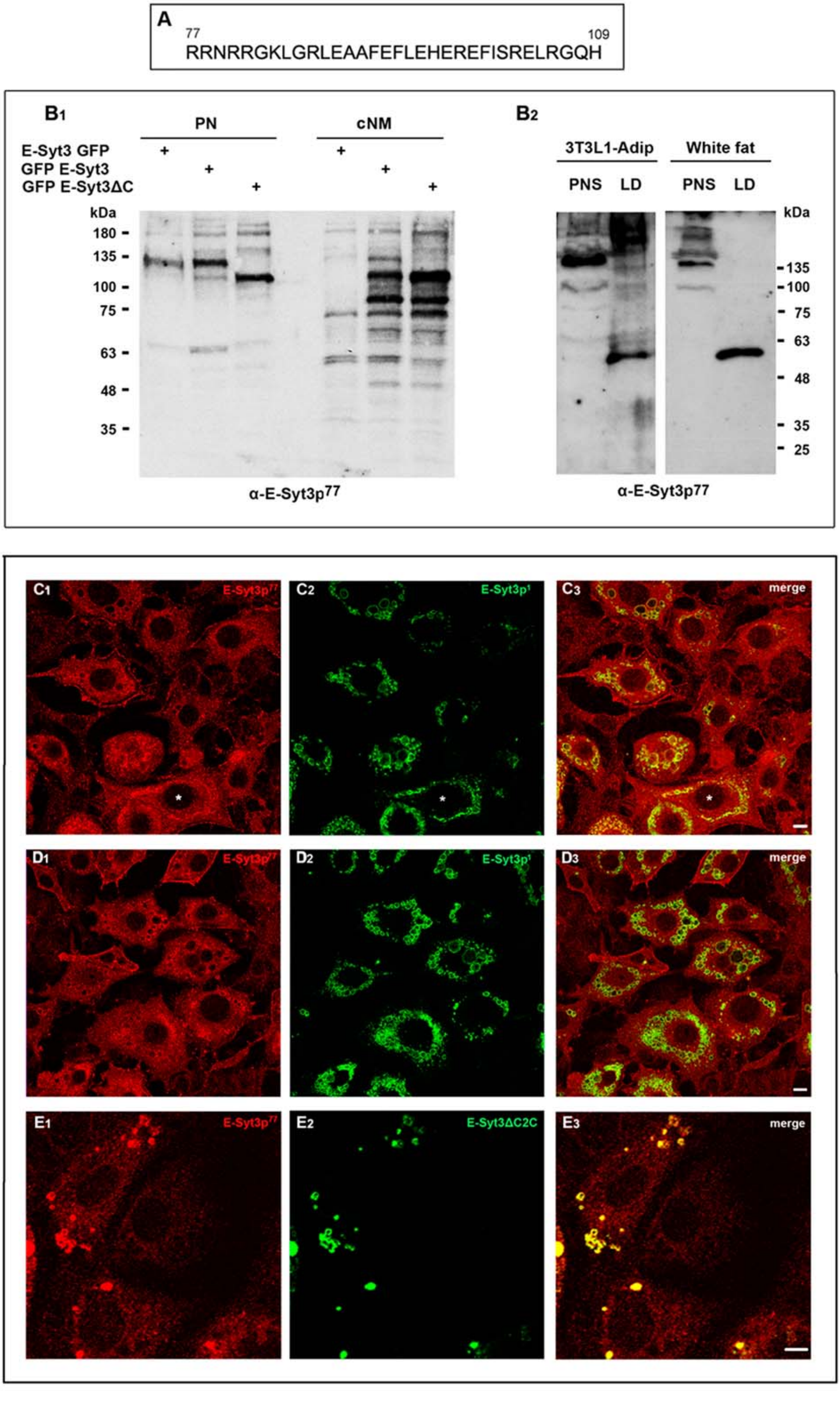

**Fig. S3 Validation of rat α-E-Syt3p$^{77}$ antibody.**

(A) The polyclonal α-E-Syt3p$^{77}$ antibody was raised in rat against the amino acid sequence $^{77}$RRNRRGKLGRLEAAFEFLEHEREFISRELRGQH$^{109}$ immediately adjacent to the hydrophobic N-hairpin that anchors the protein to the ER membrane. Partial modeling of the domain containing the immunogenic peptide using data provided by E-Syt2 and mitochondrial lipid transporter Mmm1 (PDB, ID 5YK6) suggested the reaction of α-E-Syt3p$^{77}$ with the surface of an α-helix. ($B_1$) Western blot analysis of the α-E-Syt3p$^{77}$ reaction with transfected EGFP-E-Syt3, E-Syt3-EGFP, and EGF-E- Syt3ΔC2C and the C-cleaved E-Syt3 products in the postnuclear (PN) and crude nuclear membrane (cNM) fractions (20 µg) from 3T3-L1 adipocytes, and with the PN fraction (10 µg) and purified LDs (15 µg) from 3T3-L1 adipocytes and white adipose tissue ($B_2$). (C, D) IFM of preadipocytes and young adipocytes on day 3 of differentiation incubated with α-E-Syt3p$^{77}$ antibody. In the preadipocytes, note the strong fluorescence of the ER and PM typical of ER/PM junctions, and in young adipocytes the staining of the LDs, primordial cisterna (*), PM, and ER. Also noticeable is the stronger reaction of the antibody with small/medium-sized LDs as compared to large LDs. The differences between α-E-Syt3p$^{77}$ and α-E-Syt3p$^{141}$ antibodies indicate dissimilarities in their access and/or reactivity with the epitopes in fixed cells. (E) α-E-Syt3p$^{77}$ reaction with transfected EGF-E- Syt3ΔC2C in young 3T3-L1 adipocytes. Scale bars = 10 µm.

Appendix Fig. S4

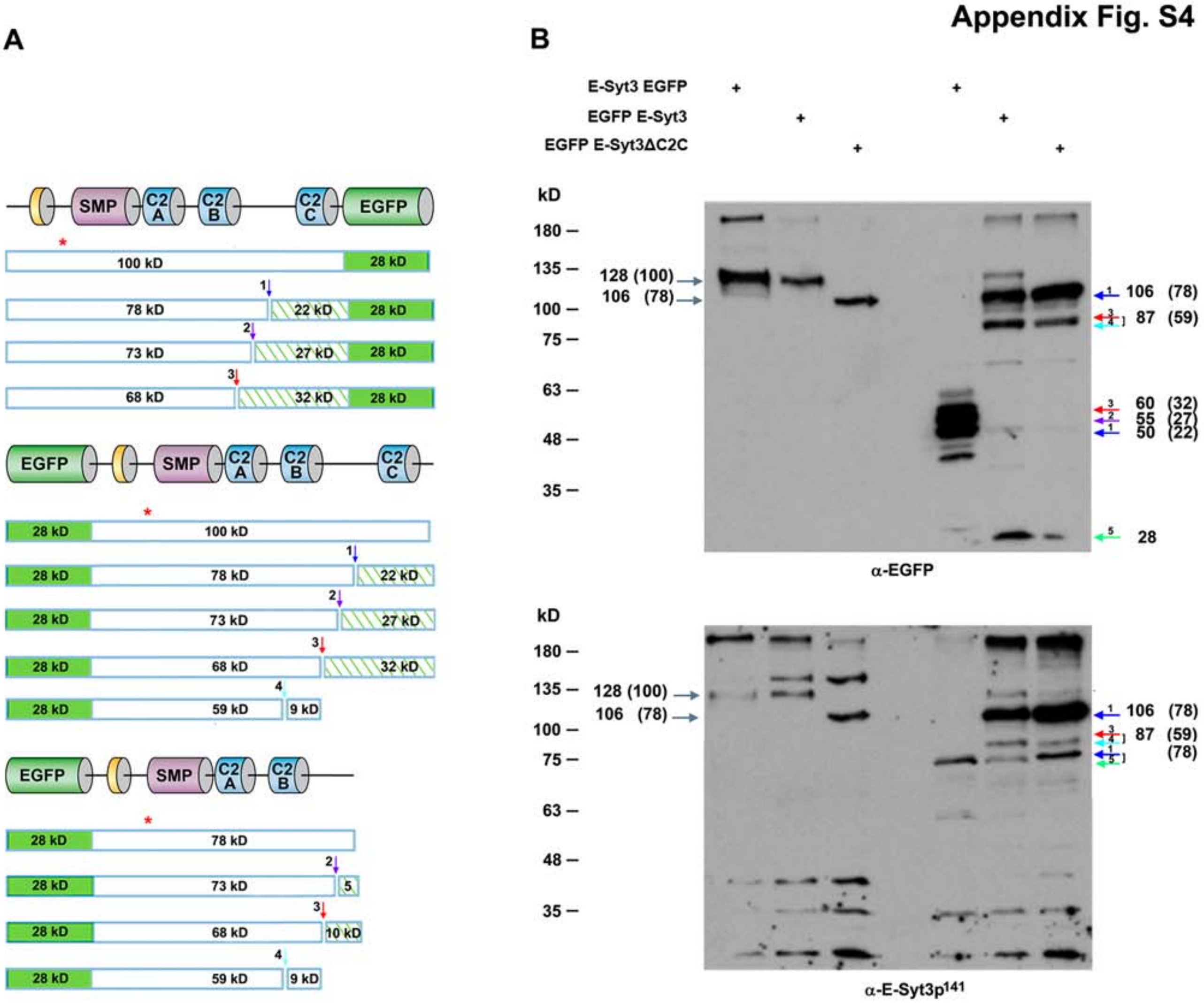


**Fig. S4 Analysis of the C-cleaved E-Syt3 fragments.**

(A) Schematic of the C-cleavage of EGFP-tagged E-Syt3 constructs. (B) Proteolytic cleavage of E-Syt3 in 3T3-L1 adipocytes separately transfected with E-Syt3-EGFP, EGFP-E-Syt3, and EGFP-E-Syt3ΔC2C (↓) using α-EGFP and α-E-Syt3p$^{141}$ antibodies (data from Fig. 2A). C-cleavage of E-Syt3 occurs at four sites (1 ↓, 2 ↓, 3 ↓, 4 ↓), with occasional removal of the 28 kDa GFP (5↓). The full-length EGFP-tagged constructs and resulting proteolytic fragments are indicated by plain numbers. The size of the full-length E-Syt3 and E-Syt3 fragments is indicated by numbers in parentheses. Sites 1, 2, and 3 are deduced from the size of the C-E-Syt3-EGFP fragments. Cleavage of E-Syt3-EGFP at site 1 produces the 50 kDa E-Syt3-EGFP and 78 kDa E-Syt3 fragments. Cleavage of EGFP-E-Syt3 at site 1 produces the 106 kDa EGFP-E-Syt3 fragment, which co-migrates with EGFPΔC2C, and a 22 kD E-Syt3 fragment. Whether the constructs are successively cleaved at more than one site by the same or different proteases (bracketed arrows) is not known. Cleavage of the 106 kDa peptide at site 4 before or after cleavage at sites 2 and 3 produces the 87 KDA EGFP-E-Syt3 fragment, whereas cleavage at site 1 before or after the removal of GFP would produce the 78 KDa E-Syt3 fragment.

Appendix Fig. S5

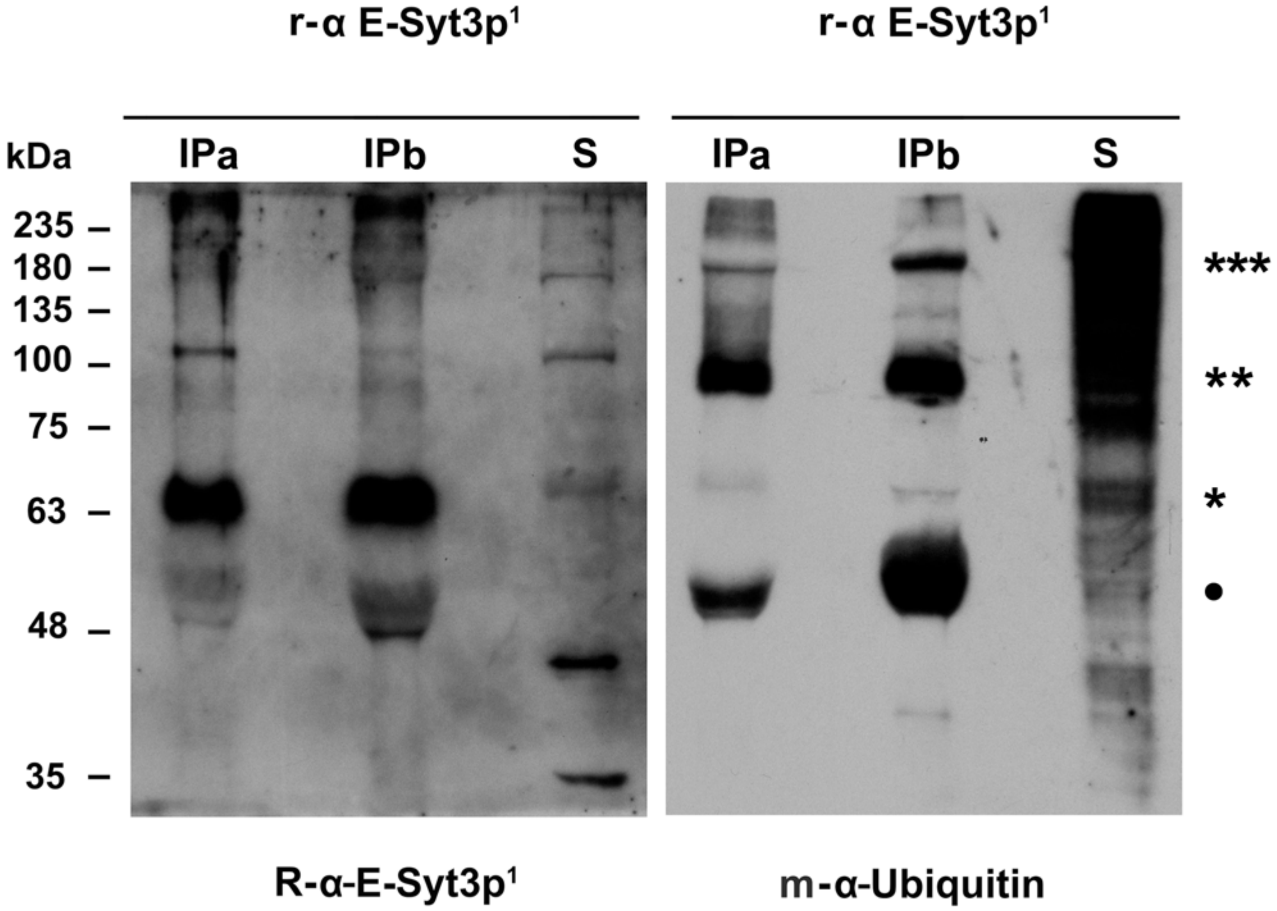


**Fig. S5 E-Syt3 ubiquitination.**

On day 4 of differentiation, 3T3-L1 adipocytes were treated with Triton X-100 and after removal of insoluble material, 200 (a) and 400 µg (b) of the supernatant were incubated for 3 h at 4°C with rat (r) α-E-Syt3p[1] Protein G Sepharose. Immunoprecipitates (IP) and supernatants (S) were subjected to western E-Syt3 ubiquitination analysis using rabbit α-E-Syt3p[1] and mouse (m) α-ubiquitin antibodies. Note that the major truncated E-Syt3 species (*) is hardly ubiquitinated whereas the minor 90 kD (**) and 180 kD (***) species produced strong ubiquitin signals (***). It is uncertain whether the 50 kD protein (●) is an ubiquitinated degradation product or a C-cleaved fragment. The experiment was repeated twice with comparable results.

**Appendix Fig. S6**

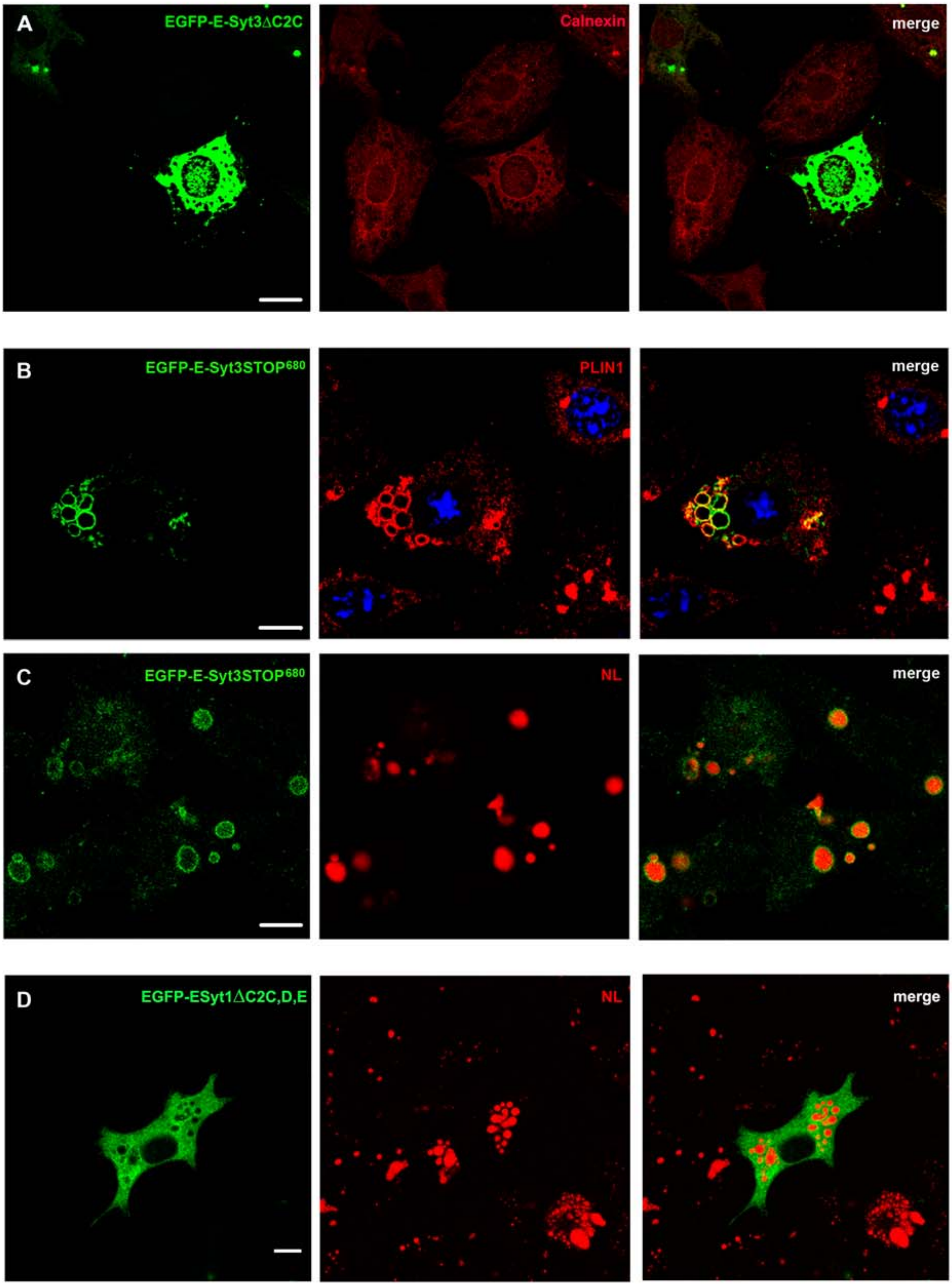

**Fig. S6 Cellular localization of truncated E-Syt3 and E-Syt1 constructs.**

(A) Overexpressed EGFP-E-Syt3ΔC2C is confined to the calnexin-positive ER of 3T3-L1 fibroblasts. Note the absence of the surface punctate staining characteristic of ER/PM junctions. (B, C) Association of EGFP-E- Syt3STOP$^{680}$ with LDs. Note that removal of the 35 aa C-tail neosequence carried by EGFP-E-Syt3ΔC2C does not affect the targeting of the truncated protein to the surface of PLIN1/NL-positive LDs. (D) Truncated E-Syt1ΔC2C,D,E is not targeted to LDs. Scale bars = 10 µm. All experiments were repeated at least twice on duplicate samples with the same result.

**Appendix Fig. S7**

**$A_1$**

$^{1}$MEHSPEEGASPEPSGQ$^{16}$
$^{1005}$DKNRSTKRKTPQKKRT$^{1020}$
$^{1038}$TLRRKLDVSVKSNSS$^{1052}$

**$A_2$**

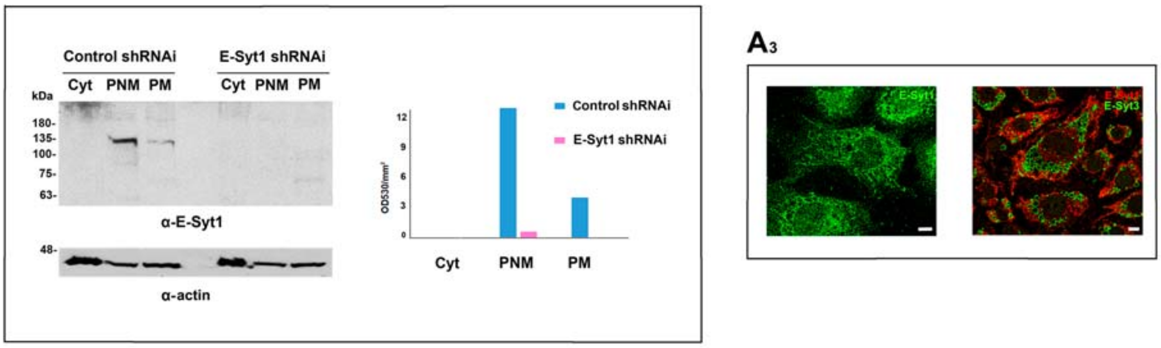


**Fig. S7 Validation of rat α-E-Syt1 antibody.**

($A_1$) Rabbit polyclonal α-E-Syt1 antibody was raised against the peptides C-$^{1}$MEHSPEEGASPEPSGQ$^{16}$, C-$^{1038}$TLRRKLDVSVKSN$^{1052}$, and C-$^{1038}$TLRRKLDVSVKSNSS$^{1020}$. ($A_2$) Western blot study of α-E-Syt1 reactivity with cytosol (CYT, 15 µg), PNM (15 µg), and purified plasma membrane sheets (PM, 15 µg) from control 3T3-L1 adipocytes and 3T3-L1 adipocytes subjected to E-Syt1 shRNA interference. Actin was used as a loading control. Note the specific reaction of the antibody with the 110 kDa E-Syt1, and the E-Syt1 repression by shRNA interference. ($A_3$) Confocal IFM. 3T3-L1-fibroblasts (left) and 3T3-L1 adipocytes (right) stained using the anti-E-Syt1 and anti-E-Syt3p[1] antibodies. Observe the localization of E-Syt1 in the ER of fibroblasts and adipocytes and the localization of E-Syt3 in the LDs. Scale bars = 10 µm.

Appendix Fig. S8

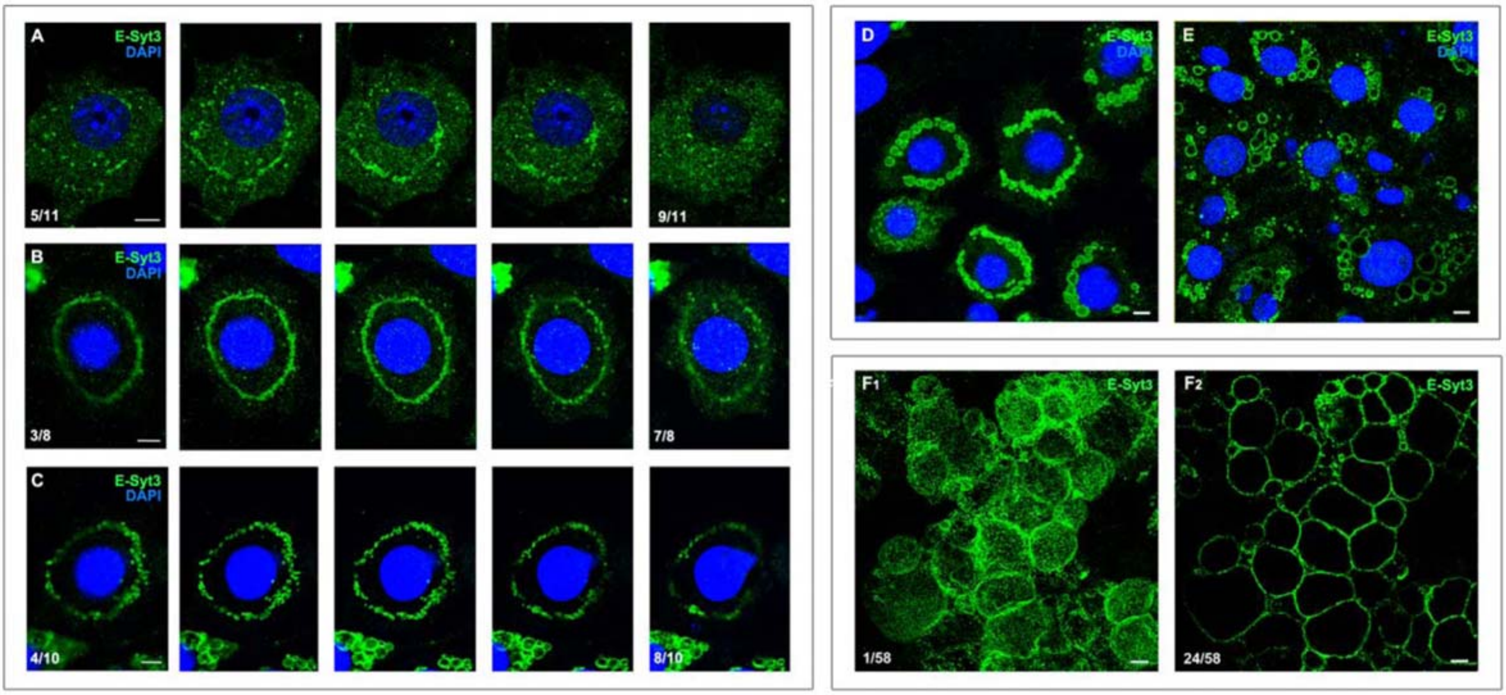


**Fig. S8 Grown LDs retain the circular arrangement of the primordial cisterna before fusing into a single giant LD.**

Confocal microscopy of the primordial cisterna (A-C) and LDs (D-G) in 3T3-L1 adipocytes on days 3 (A -C), 4 (D, E), and 24 ($F_{1-2}$) of differentiation using antibody α-E-Syt3p[141]. (A-C) 1 µm consecutive sections; the average cisterna measured 74±9.5-µm-long and 2-µm-thick; the longest was 147 µm and the shortest 40 µm (n= 10). (D, E) Observe the growth and formation of 63.5±5.4 µm long LD collars (n=10) that retained the shape of the primordial cisterna, and the assembly of single-giant LDs in 24-day-old 3T3-L1 adipocytes ($F_{1, 2}$). Scale bars = 5 µm. Bottom left numbers indicate the section position within the stack.

### Appendix Fig. S9

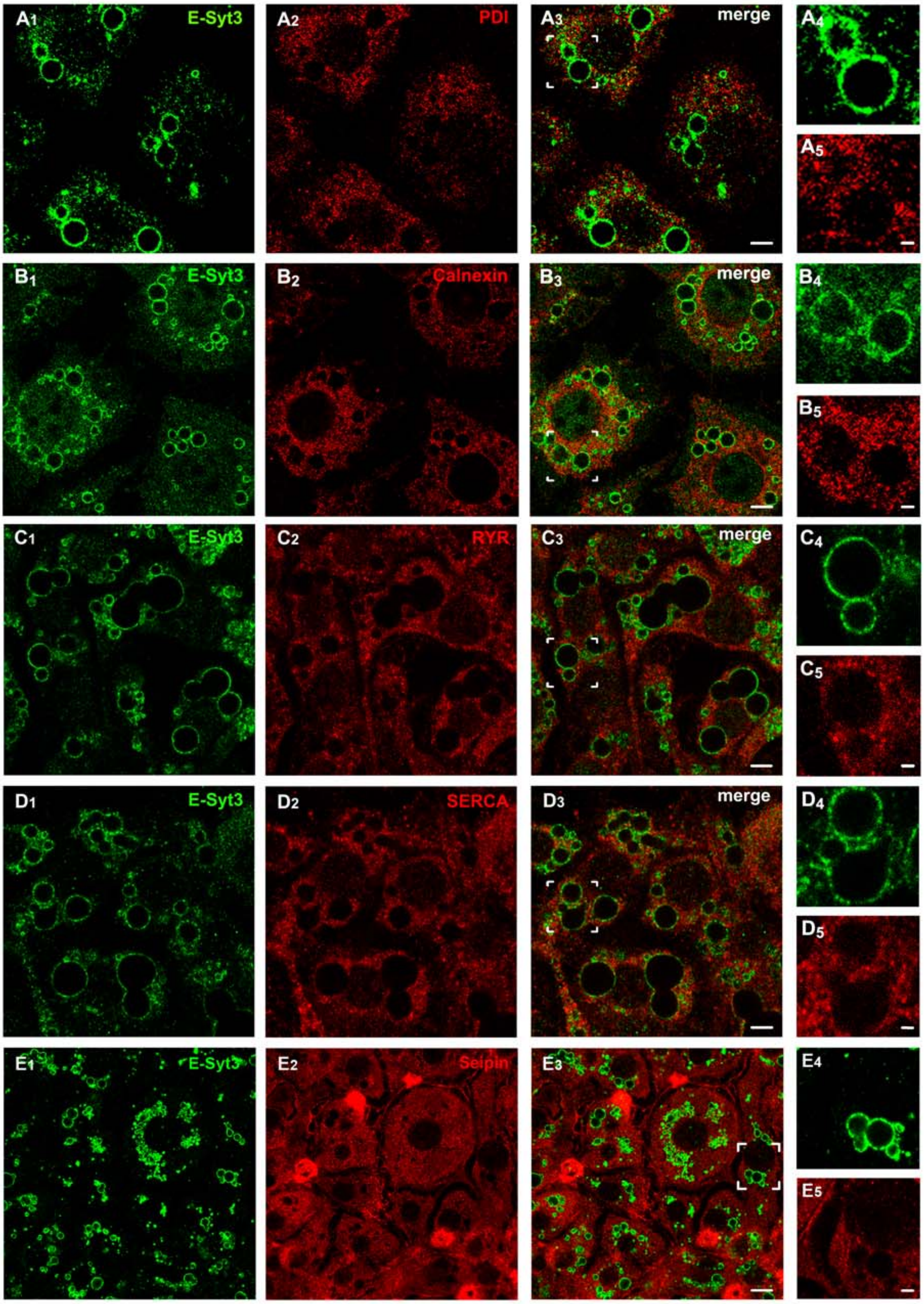

**Fig. S9 ER membrane protein markers do not coat grown LDs.**

Note the encapsulation of LDs within the sharp E-Syt3 boundaries from which the ER protein markers protein disulfide isomerase (PDI), calnexin, ryanodine receptor (RYR), sarco/endoplasmic reticulum $Ca^{2+}$-ATPase (SERCA), and seipin are absent, as well as the spreading of the markers throughout the extensive ER. Scale bars = 10 µm (panels 1-3) or 2.5 µm (panels 4 and 5).

#### Appendix Fig.S10

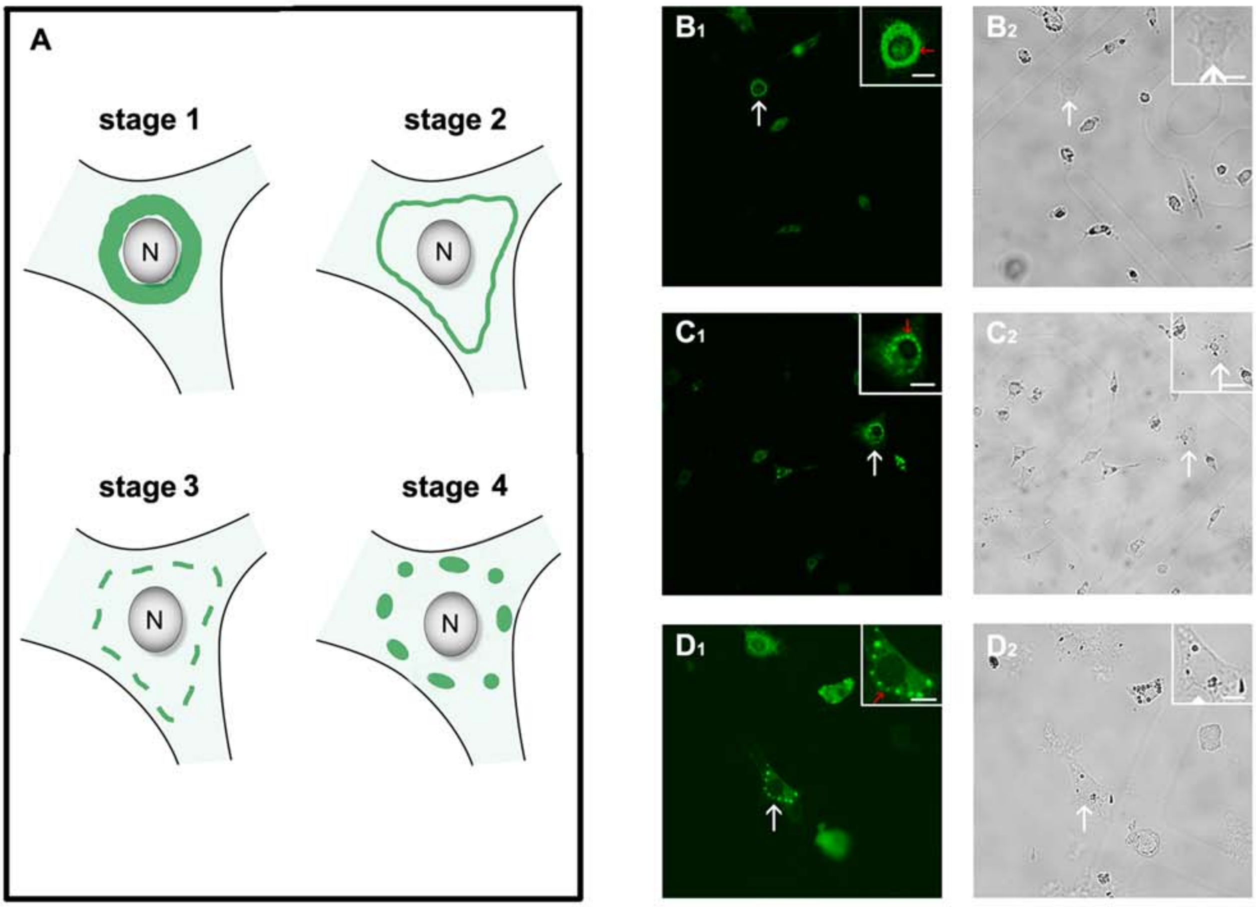


**Fig. S10 Correlative IFM-bright field microscopy (CLEM) for adipocyte selection to study LD biogenesis by EM and TM.**

(A) Schematic describing the changes in EGFP-E-Syt3ΔC2C distribution during relocation from the early synthesis to primordial cisterna and LDs. The four main stages description is based on the experimental data shown in Figs. 4 and S8A-C ,and videos 1 and 2. Stage 1, broad and circular arrangement of the protein in the cytoplasm before becoming concentrated in the primordial cisterna; stage 2, concentration of the protein in the primordial cisterna; stage 3, segmentation of the cisterna; stage 4, concentration of the protein into globular cisterna segments. (B, D) Selection of preadipocytes and young adipocytes for high resolution EM ultrastructural studies. On day 3 of differentiation, the cells were transfected for 6 h with EGFP-E-Syt3ΔC2C; white arrows mark preadipocytes at stage 1 ($B_1$, $B_2$) and young adipocytes at stage 4 ($C_{1-2}$, $D_{1-2}$) (see Fig. 5). Note in the inserts (C, D) the clear distinction between the dark LDs and the fluorescent cisterna segments. Scale bars = 40 µm.
